# The BAF complex works with FOS to regulate human neuronal activity-dependent ASD-associated gene programs

**DOI:** 10.64898/2026.08.25.747063

**Authors:** Sara K. Trowbridge, GiHun Choi, Ava C. Carter, Jillian E. Petrocelli, Josephine E. Robb, Gabriel T. Koreman, Siwei Chen, Ryan N. Doan, Christopher P. Davis, David A. Harmin, Eric C. Griffith, Konrad J. Karczewski, J. Wade Harper, Michael E. Greenberg

**Affiliations:** Department of Neurobiology, Harvard Medical School, Boston, MA, 02115 USA; Allen Discovery Center for Human Brain Evolution, Boston, MA, 02115, USA; Hock E. Tan and K. Lisa Yang Center for Autism Research at Harvard University, Boston, MA, 02115, USA; Blavatnik Institute, Harvard Medical School, Boston, MA, 02115, USA; Department of Neurology, Boston Children’s Hospital, Boston, MA, 02115, USA; Analytic and Translational Genetics Unit, Massachusetts General Hospital, Boston, MA, 02114, USA; Program in Medical and Population Genetics, Broad Institute of MIT and Harvard, Cambridge, MA, 02142, USA; Novo Nordisk Foundation Center for Genomic Mechanisms of Disease, Broad Institute of MIT and Harvard, Cambridge, MA, 02142, USA; Division of Genetics and Genomics, Boston Children’s Hospital, Boston, MA, 02115, USA; Department of Cell Biology, Blavatnik Institute, Harvard Medical School, Boston, MA, 02115 USA

**Author notes:** Corresponding author: Michael E. Greenberg. These authors contributed equally (Sara K. Trowbridge, GiHun Choi, Ava C. Carter).

## Abstract

The BAF chromatin remodeling complex is critical to normal brain development, and rare variants within genes encoding BAF subunits are a common genetic cause of neurodevelopmental disorders, including autism spectrum disorder (ASD). Yet the factors that direct BAF binding across the neuronal genome, the human neuronal gene programs that are regulated by BAF, and the mechanisms whereby BAF subunit perturbation leads to ASD are not known. We find that BAF binds with the activity-dependent transcription factor FOS to distal regulatory regions that, in response to neuronal activity, undergo chromatin opening and show evidence of enhancer activation. Knock-out of ARID1A, a BAF subunit implicated in ASD, leads to decreased chromatin accessibility at FOS/BAF binding sites concomitant with decreased expression of nearby activity-regulated ASD-associated genes. Additionally, we find that the FOS binding motif in FOS/BAF-bound regions is highly constrained in the human population, and that rare variants in this motif in ASD-affected individuals disrupt stimulus-dependent enhancer activation. This suggests that genetic variation in FOS/BAF-bound regions contributes to ASD pathogenesis, due to an inability to recruit FOS and BAF to enhancers to promote gene expression. Together, our findings highlight a role for BAF in mediating neuronal transcriptional programs downstream of FOS and reveal a mechanism by which non-coding variants may impact BAF function and contribute to risk for ASD.

## Introduction

Rare variants within genes that encode chromatin regulators are a common genetic cause of autism spectrum disorder (ASD), underscoring the importance of the tight regulation of chromatin state and gene transcription for brain development and function^1^. The BRG1/BRM-associated factor (BAF) complex is a large chromatin remodeling complex comprised of at least 15 subunits encoded by 29 different genes, nearly half of which have been implicated in ASD or other neurodevelopmental disorders^2,3^. BAF function has been studied in detail in neural progenitor cells and in the developing nervous system, and BAF has been shown to be critical for developmental processes, such as neural stem cell proliferation, neuronal differentiation, and neuronal migration^2^. However, in post-mitotic neurons, the factors which guide BAF to specific regions of the genome and the direct BAF transcriptional targets were previously unidentified. Given evidence that mutations within BAF subunits lead to significant neurodevelopmental abnormalities in humans, identifying the sites of BAF binding, the direct gene targets regulated by BAF, and overall BAF function in human neurons is of particular interest.

In this study we investigated the possibility that BAF regulates activity-dependent gene expression in human neurons. In neurons, membrane depolarization in response to neurotransmitter release at synapses leads to calcium influx and activation of signaling cascades that trigger the rapid transcription of immediate early gene (IEG) transcription factors (TFs), including the AP-1 family member FOS^4^. These IEGs then activate late response genes (LRGs), which encode effector proteins, including synaptic and secreted molecules that mediate aspects of neuronal maturation and plasticity^5,6^. While activity-dependent neuronal gene programs are critical for nervous system development and function throughout life, whether and how BAF contributes to these gene programs remained to be determined.

Recent work from our laboratory demonstrated that in fibroblasts BAF interacts with the AP-1 TF FOS^7^, suggesting that FOS might mediate BAF binding across the neuronal genome and that BAF is a critical regulator of neuronal activity-dependent transcription downstream of FOS. However, since BAF incorporates neuronal-specific subunits which change over the course of neural development^8–10^, it was possible that, unlike in fibroblasts, BAF functions independently of FOS in post-mitotic neurons.

We also considered the possibility that the study of BAF and FOS signaling in human neurons might provide insight into the contribution of non-coding variation to human disease, an especially challenging topic due to the difficulty of predicting the functional effects of noncoding variants^11^. Previous studies have identified a significant enrichment of single nucleotide polymorphisms (SNPs) associated with common neuropsychiatric disorders within the gene regulatory elements that mediate activity-dependent gene transcription^12,13^. However, it was not known which SNPs are causal for disease, nor how SNPs disrupt activity-dependent TF binding and gene expression. We hypothesized that, if FOS and BAF co-regulate neuronal gene programs, sequence variants that disrupt FOS binding could also impact BAF-regulated programs important for neurodevelopment.

Here, using human embryonic stem cell (hESC)-derived neurons^14^, we find that FOS and BAF co-bind putative enhancer regions in an activity-dependent manner. These FOS/BAF binding sites are near many activity-induced ASD-associated genes, a subset of which display decreased expression when ARID1A, a core BAF complex subunit implicated in ASD^15,16^, is disrupted, suggesting a possible mechanism whereby mutations in BAF subunits cause ASD.

We also find that AP-1 motifs (TGASATCA) within neuronal FOS/BAF-bound regions are conserved across evolution and are highly constrained in humans, indicating that rare variants within these regulatory regions would be likely to disrupt enhancer function and affect nearby gene expression. Consistent with this, we demonstrate that rare ASD patient variants in AP-1 motifs of several FOS/BAF binding sites near ASD-associated genes disrupt activity-dependent enhancer function, highlighting one mechanism by which rare variants in non-coding regions may perturb gene expression and subsequent nervous system development.

## Results

### Identification of human neuronal activity-dependent FOS binding sites

Human neuronal activity-dependent gene programs have been challenging to study using available post-mortem or surgical samples because the expression of activity-dependent IEGs is tightly time-locked to stimulus, and most neurons in unstimulated human brain samples do not express IEGs. The analysis of activity-dependent IEG TF binding in the human brain is further complicated by cell type heterogeneity, as IEG TFs bind to the genome in a highly cell-type-specific manner^7,17,18^. We therefore employed a relatively homogeneous population of human neurons that could be synchronously exposed to stimuli that lead to the robust expression of IEGs, thus providing a neuronal population with which to study BAF-regulated activity-dependent programs.

Using an engineered H9 hESC line harboring a doxycycline-inducible NGN2 cassette within the *AAVS1* safe-harbor locus^19^, we adapted a previously described hESC differentiation protocol combining inducible pro-neurogenic *NGN2* expression with dual SMAD and WNT inhibition to induce rapid patterning toward forebrain excitatory neurons (human patterned induced neurons, hpiNs; Fig. 1a)^14^. After four weeks in culture, hpiNs robustly express neuronal markers NF and MAP2 and the cortical neuronal marker BRN2, as confirmed by immunohistochemistry (Extended Data Fig. 1a).

**Figure 1.**
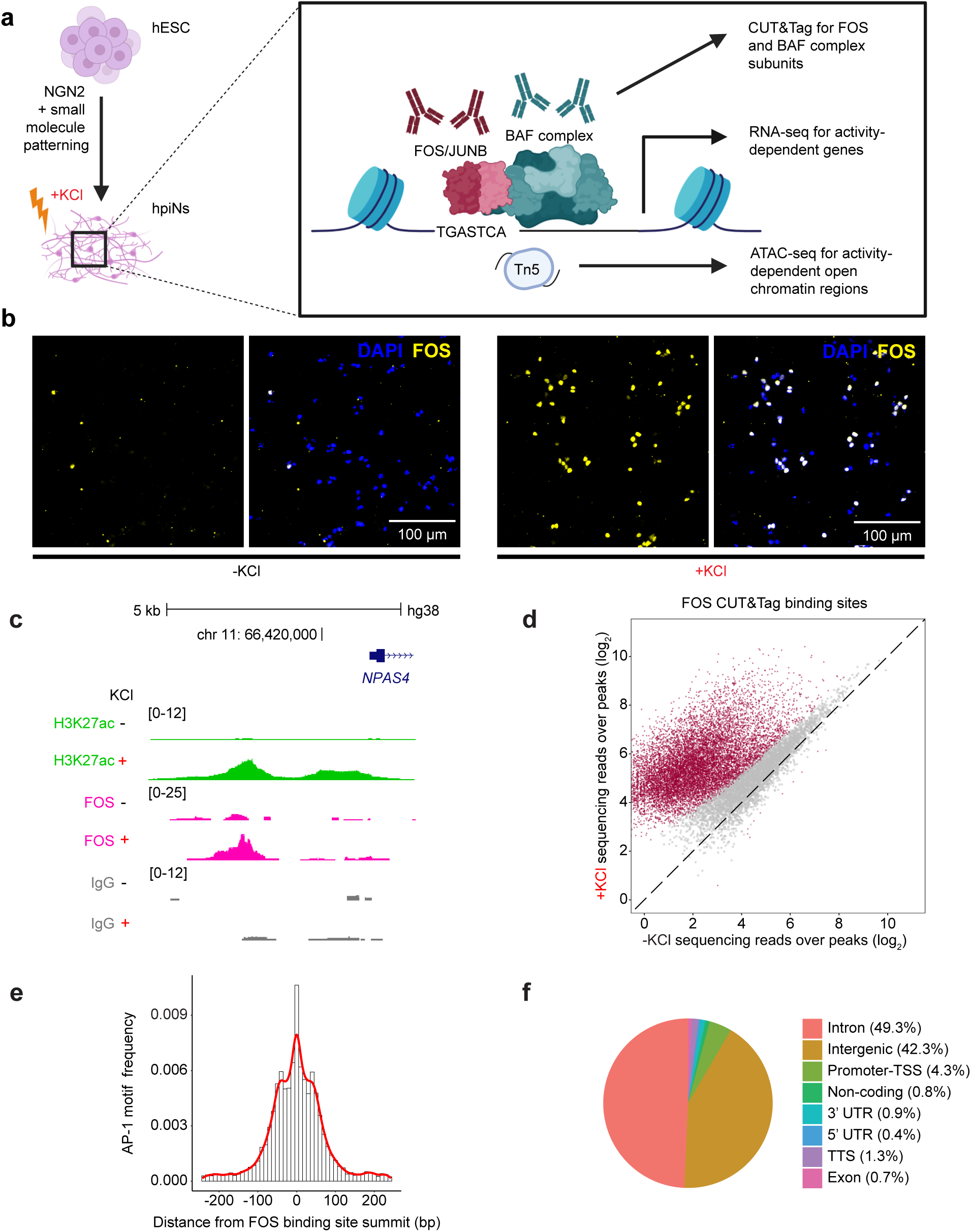
Identification and characterization of activity-dependent FOS binding sites. (**a**) Experiment schematic. (**b**) Representative images of FOS immunohistochemistry in hpiNs before (-KCl) or 2 hr after depolarization (+KCl). (**c**) Representative genome browser tracks of CUT&Tag data for FOS (n = 6, pink), H3K27ac (n = 3, green), and paired IgG negative control upstream of the *NPAS4* locus before (-) and 2 hr after depolarization (+). Scale is indicated in the brackets. (**d**) Scatter plot of CUT&Tag sequencing reads (unstimulated vs 2 hr post-depolarization) for all FOS binding sites. Sites with significantly different FOS binding between timepoints are highlighted in red (n = 6 biological replicates, adjusted p<0.05). (**e**) Distribution of the AP-1 motif relative to CUT&Tag peak signal in all FOS binding sites. (**f**) Pie chart of genomic annotations for FOS binding sites.

Prior work demonstrated that *NGN2* overexpression without small molecule patterning yields a heterogeneous population of neurons with mixed peripheral (PNS) and central nervous system (CNS) identities^20^. To characterize the extent of heterogeneity within our hpiN cultures, we performed single-cell RNA-seq (scRNA-seq) and found that the hpiNs are highly homogeneous, express CNS but not PNS markers, and express excitatory but not inhibitory neuronal markers (Extended Data Fig. 1b). Bulk RNA-seq data confirmed expression of upper layer cortical markers *BRN2* and *CUX1*, with lower expression of lower layer markers *CTIP2* and *TBR1*. Similar findings were obtained for hpiNs derived from two additional hESC lines, H1 and HUES64 (Extended Data Fig. 1c). Overall, this suggests that hpiNs are most similar to an excitatory upper layer cortical neuronal population.

To trigger synchronized neuronal depolarization, we exposed hpiNs to elevated levels of potassium chloride (KCl), a robust stimulation protocol that allows for the identification of physiologically-relevant activity-dependent gene programs *in vitro*^21,22^. *FOS* expression is elevated after treatment with 55 mM KCl but is induced to a lesser extent in the presence of the L-type voltage-gated calcium channel blocker nimodipine or the calcium chelator EDTA, indicating that calcium entry into the neurons is critical for *FOS* induction (Extended Data Fig. 1d). By scRNA-seq and immunohistochemistry, we confirmed the widespread induction of FOS mRNA and protein within 2 hours (hr) of KCl exposure (Fig. 1b, Extended Data Fig. 1b,e).

To determine if BAF is recruited to FOS binding sites in neurons, we first assessed FOS binding across the neuronal genome using CUT&Tag^23^. Comparison of FOS CUT&Tag signal between the unstimulated and 2 hr KCl-stimulated H9-derived hpiNs revealed 8,728 binding sites that display increased FOS binding after membrane depolarization and contain at least one canonical AP-1 motif (TGASTCA) within the FOS binding site (Fig. 1c-e). The FOS CUT&Tag signal was abolished over these FOS binding sites in FOS knock-out hpiNs (Extended Data Fig. 2a-b), confirming the specificity of the antibody and protocol. Consistent with reported FOS binding patterns in rodent neurons^17^, the vast majority of FOS binding sites are located within intronic or intergenic regions (7,994/8,728, 92%, Fig. 1f). Many of these FOS binding sites (6,790/8,728, 78%) overlap enhancers identified *in vivo*^24,25^ from primary human brain tissue and are reproducible across hpiNs derived from two other hESC lines (H1 and HUES64) (Extended Data Fig. 2c-d). For on-going analyses, we focused on the 8,728 binding sites identified in H9-derived hpiNs as a high-confidence set of activity-dependent FOS binding sites in human excitatory neurons (Supplementary Table 3).

### FOS and BAF co-bind activity-responsive gene regulatory elements near ASD-associated genes

We next asked if BAF binds human neuronal activity-dependent FOS binding sites. Interactions between BAF and other AP-1 factors have been observed in neural progenitor cells and non-neuronal cell types^7,26–28^, but were not previously characterized in post-mitotic neurons. We performed CUT&Tag for four BAF complex subunits (ARID1A, ARID1B, SMARCB1 and SMARCC1) in unstimulated and membrane-depolarized hpiNs. These BAF subunits were selected due to their known roles in mediating the binding of BAF to DNA^29^. The CUT&Tag signal for all four BAF subunits is significantly increased at FOS binding sites 2 hr after membrane depolarization (p<2.2e-16, n=3) (Fig. 2a-b), a finding that is reproducible across hpiNs derived from the H1 and HUES64 lines (Extended Data Fig. 3a). RNA expression of these subunits is not increased after membrane depolarization, suggesting that the increased binding at FOS-bound sites is due to recruitment of pre-existing protein.

**Figure 2.**
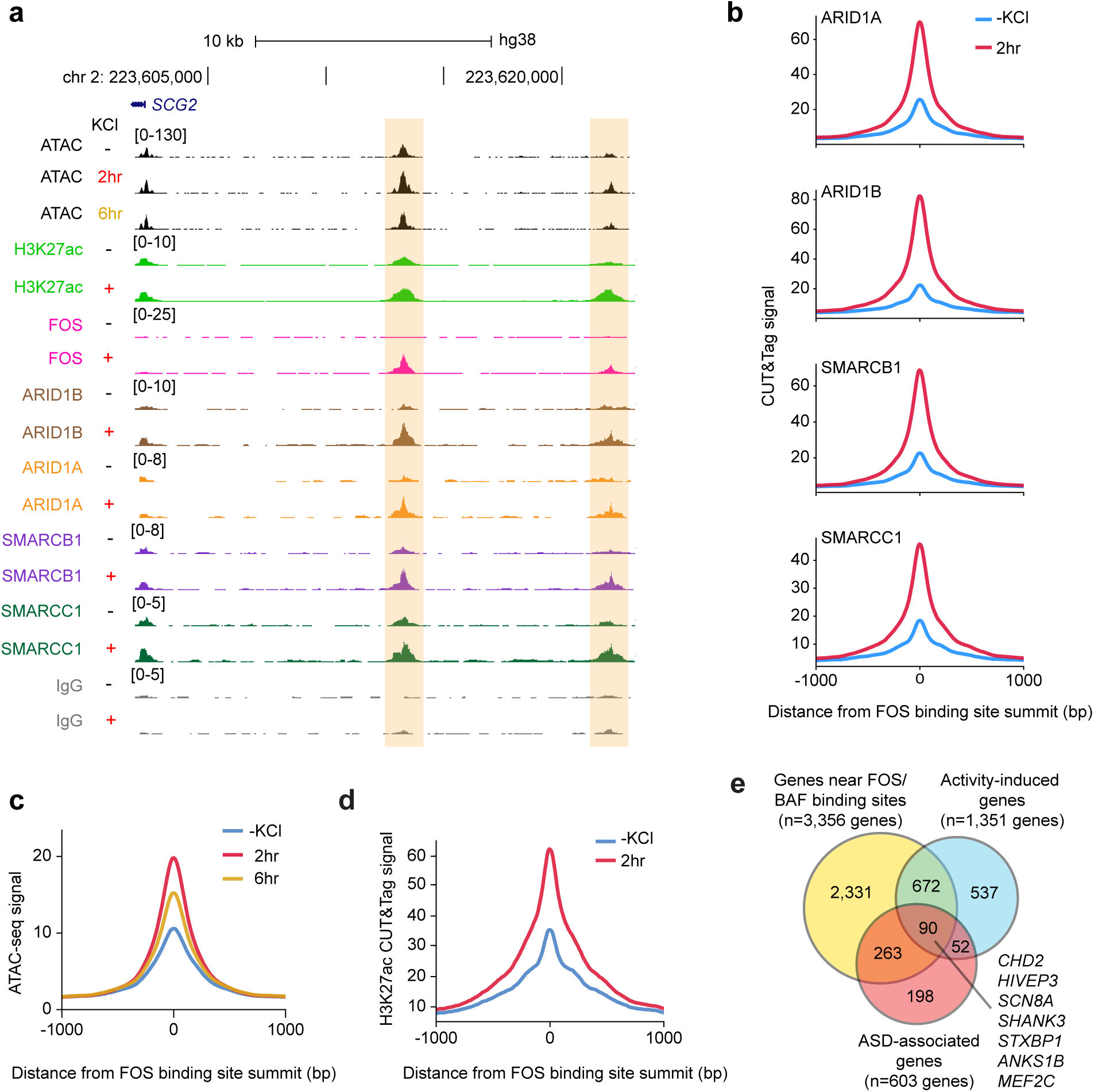
The BAF complex binds FOS binding sites in an activity-dependent manner. (**a**) Representative genome browser tracks of ATAC-seq data (n = 3, black) and CUT&Tag data for H3K27ac (n = 3), FOS (n = 6), BAF complex subunits ARID1B, ARID1A, SMARCB1, and SMARCC1 (all n = 3), and paired IgG negative control near the *SCG2* locus before (-) or 2 hr after (+) depolarization. Scale is indicated in the brackets. (**b**) Aggregate enrichment plots of BAF complex subunit CUT&Tag signal from hpiNs before (blue) or 2 hr after (red) depolarization over all FOS binding sites. (**c**) Aggregate enrichment plot of ATAC-seq signal over FOS/BAF binding sites before (blue), 2 hr (red), or 6 hr after (yellow) depolarization (n = 3). (**d**) Aggregate enrichment plot of H3K27ac CUT&Tag signal over all FOS/BAF binding sites, before (teal) or 2 hr after (red) depolarization. (**e**) Venn diagram demonstrating overlap between genes near FOS/BAF binding sites, activity-induced genes, and ASD-associated genes.

BAF remodels chromatin in an ATP-dependent manner by evicting nucleosomes to increase the accessibility of TF binding sites^30–32^. To determine whether chromatin accessibility is increased at the FOS/BAF binding sites in response to neuronal depolarization, we performed ATAC-seq (assay for transposase-accessible chromatin using sequencing) at three timepoints (-KCl, 2 hr, and 6 hr) after neuronal depolarization. At FOS/BAF binding sites, chromatin accessibility is significantly increased 2 hr after membrane depolarization and then decreases towards baseline after 6 hr of KCl exposure (Fig. 2c). This increased accessibility may be due to the ability of BAF to remodel nucleosomes by evicting histones, rendering the regulatory elements accessible for stable binding of FOS and other factors. Additionally, we observed an activity-dependent increase in the active enhancer signal H3K27ac (Fig. 2d), suggesting that FOS/BAF binding promotes enhancer activation.

To test whether activity-dependent BAF recruitment is a specific feature of FOS-bound regions, we defined a separate set of activity-dependent binding sites for the TF NPAS4, a highly inducible IEG that interacts with the chromatin remodeler NuA4 but not the BAF complex^33^ (Supplementary Table 3). We found that inducible BAF subunit binding is significantly less at NPAS4 binding sites as compared to FOS binding sites (Extended Data Fig. 3b-d). Additionally, we identified open chromatin ATAC-seq regions with high versus low inducible FOS CUT&Tag signal but comparable inducible ATAC-seq signal (n = 264 per group). Regions with high inducible binding of FOS display significantly more inducible BAF complex subunit binding as compared to regions with low inducible binding of FOS (Extended Data Fig. 3e-f). This indicates that inducible BAF binding is not a general feature of activity-responsive regions but instead specifically correlates with activity-dependent FOS binding. Along with prior evidence that FOS and BAF can be co-precipitated from fibroblast extracts^7^, these analyses suggest a model whereby FOS recruits the BAF complex to nucleosome-bound enhancers that contain AP-1 sites, and BAF evicts nucleosomes and increases chromatin accessibility, leading to enhancer activation.

Cell-type-specific TFs have been hypothesized to function in concert with FOS to specify cell-type-specific, stimulus-dependent gene programs^7^. To identify such cell-type-specific factors that function in hpiNs together with the FOS/BAF complex, we scanned the FOS/BAF binding sites for consensus binding motifs for TFs other than AP-1 TFs that are expressed in hpiNs. Using this strategy, we found an enrichment for the CUX1/2 motif, as has been previously reported for FOS binding sites^13,34^, and an enrichment for the binding motif for the EBF family of TFs. Notably, 56% of FOS/BAF-bound sites contain at least one EBF binding motif (Extended Data Fig. 3g). Three of the four members of the EBF TF family are expressed in hpiNs (EBF1, 2 and 3), consistent with studies in animal models that demonstrated EBF TFs are highly expressed in the developing nervous system and important in neuronal differentiation^35,36^. Interestingly, pathogenic variants in *EBF3* cause neurodevelopmental disorders, including ASD^37,38^. The role of the EBF TFs in binding and regulation of FOS/BAF-bound regions remains unclear, but this finding raises the intriguing hypothesis that dysregulated FOS/BAF signaling is also a component of EBF3-related ASD.

To characterize the activity-dependent gene program likely to be regulated by the FOS-BAF complex, we performed bulk RNA-seq with unstimulated cultures and cultures harvested 15 minutes (min), 1, 2, or 6 hr after exposure to KCl. This analysis identified a total of 1,351 activity-induced genes (adj p < 0.05, FPKM ≥ 2, n=3, Extended Data Fig. 4a). Along with *FOS*, other canonical IEGs, including *NPAS4*, *EGR1*, and *NR4A1*, are elevated at the 15 min, 1 hr, and 2 hr time points. At 6 hr post-stimulation, we observed a wave of LRG induction, including LRGs previously characterized in mouse neurons (e.g. *SCG2*, *HDAC9* and *NFIL3*)^17,39^. These findings are consistent across H1- and HUES64-derived hpiNs (Extended Data Fig. 4b).

We next identified the expressed genes closest to each FOS/BAF-bound site (FPKM ≥ 2, n = 3356) as the genes most likely to be regulated by FOS and BAF^40^ (Supplementary Table 3). We found that more than half of activity-induced genes are near at least one FOS/BAF binding site (n = 762, 56% of all activity-induced genes). To determine whether these putative FOS/BAF target genes have a known association with ASD, we searched the SFARI Gene database^41^ and found that of the 603 SFARI genes expressed in hpiNs (FPKM ≥ 2), 353 (59%) were located near at least one FOS/BAF binding site. Overall, genes near FOS/BAF binding sites were enriched for SFARI genes as compared to expressed genes not near a FOS/BAF binding site (353/3356, 10.5% vs. 250/4540, 5.5%, Pearson’s chi-squared test p < 2.2e-16). Ninety (25%) of the SFARI genes near a FOS/BAF binding site are induced by membrane depolarization, including several well-characterized ASD-associated genes (Fig. 2e; e.g., *MEF2C, SHANK3, STXBP1, CHD2*)^42–45^. These findings demonstrate that in human neurons the FOS/BAF complex is likely to regulate genes known to be critical for normal neurodevelopment and causal for ASD.

### ARID1A knockdown decreases chromatin accessibility at a subset of FOS/BAF binding sites

Having demonstrated that FOS and BAF co-bind activity-induced regions across the genome, we next asked if disruption of the BAF complex affects activity-dependent gene transcription. We chose to focus on ARID1A, a core subunit of the canonical BAF complex, as heterozygous pathogenic variants in *ARID1A* are associated with a severe NDD, often with autistic features^15,16^. We generated hESC lines with heterozygous (ARID1A het) and bi-allelic (ARID1A KO) truncating variants in *ARID1A* and confirmed that ARID1A protein levels are reduced ∼50% and 90%, respectively (Fig. 3a-b). The ARID1A mutant cell lines differentiate into cells that have neuronal morphology and express the neuronal markers BRN2 and MAP2, as determined by immunohistochemistry (Extended Data Fig. 5a). Moreover, RNA-seq experiments demonstrated overall preserved expression of a wide range of marker genes^14^ in the ARID1A het and KO hpiNs as compared to wildtype (Extended Data Fig. 5b).

**Figure 3.**
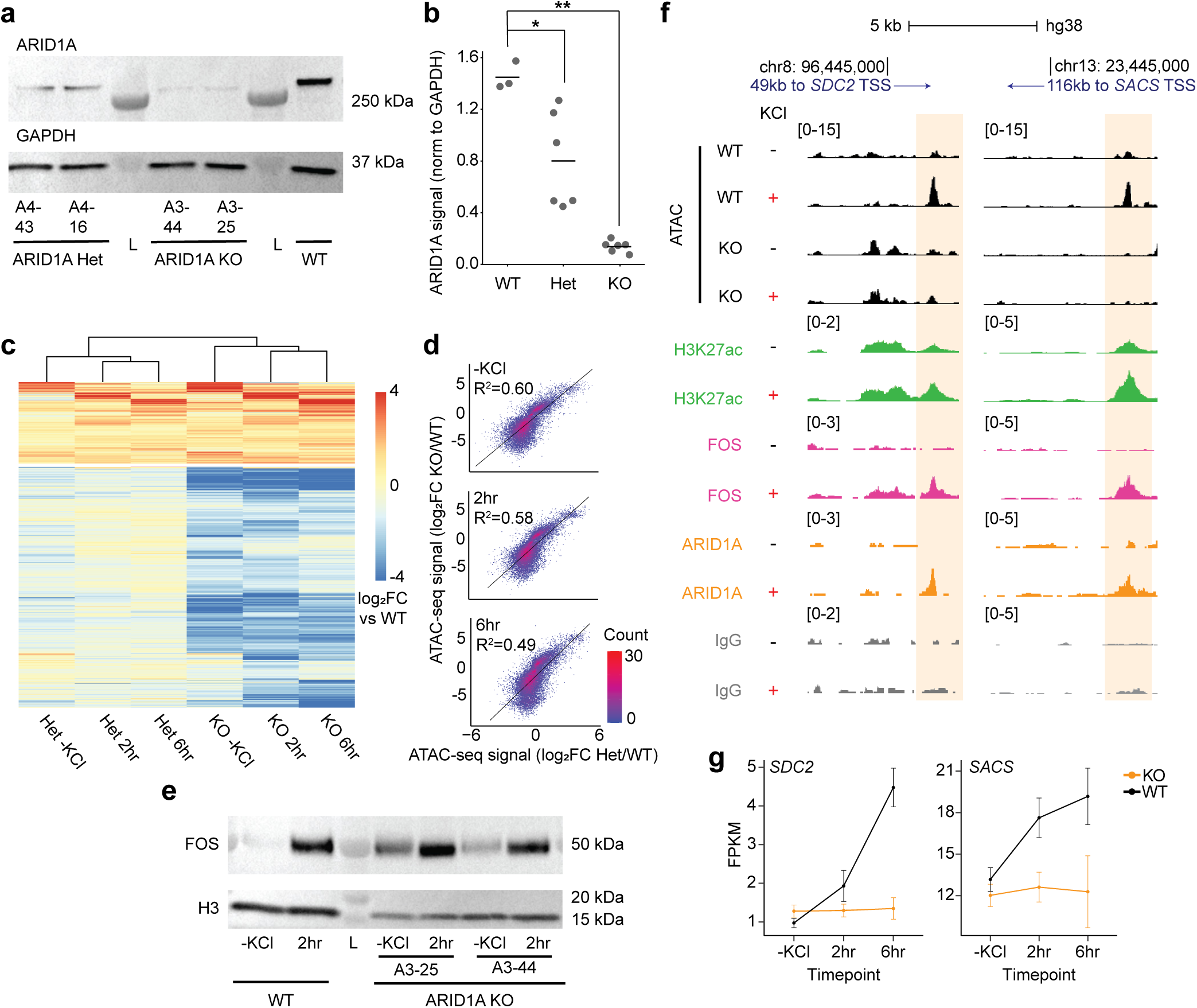
ARID1A KO leads to decreased chromatin accessibility at FOS/BAF binding sites. (**a**) Western blot of ARID1A in the ARID1A het and KO hpiNs at Day 28, along with GAPDH loading control. (**b**) Quantification of normalized ARID1A protein signal (n = 3, bar = mean, *adjusted p<0.05, **adjusted p<0.005). (**c**) Heatmap of the log2 ATAC-seq signal fold-change in the ARID1A het (Het) or ARID1A KO (KO) hpiNs versus wildtype (WT) before (-KCl), 2 hr after, or 6 hr after depolarization. Experiments were conducted with 2 ARID1A het lines with 3 biological replicates each and 1 ARID1A KO line with 3 biological replicates at 2 hr and 2 biological replicates at 6 hr, as compared to paired WT samples. (**d**) Scatter density plots showing the correlation between the log2 fold-change in ATAC signal for ARID1A het hpiNs versus wildtype and the log2 fold-change in ATAC signal for ARID1A KO hpiNs versus wildtype. (**e**) Western blot of FOS in the ARID1A KO hpiNs at Day 28, along with H3 loading control before (-KCl) or 2 hr after depolarization. (**f**) Genome browser tracks of ATAC-seq data from ARID1A KO or WT hpiNs (n = 3, black) and CUT&Tag data from WT hpiNs for H3K27ac (green, n = 3), FOS (pink, n = 6), ARID1A (yellow, n = 3), and paired IgG negative control (gray) near the *SDC2* and *SACS* loci before (-) or 2 hr after (+) depolarization. Scale is indicated in the brackets. (**g**) RNA expression (n = 3 biological replicates, mean ± s.d.) of *SDC2* and *SACS* in WT (black) or ARID1A KO (yellow) hpiNs before (-KCl), 2 hr after, or 6 hr after depolarization.

To assess the effects of decreased ARID1A expression on chromatin accessibility, we performed ATAC-seq and identified regions with differential chromatin accessibility at three timepoints (-KCl, 2 hr, and 6 hr) after membrane depolarization. Overall, 1822 regions with differential chromatin accessibility (adjusted p-value < 0.05, log_2 f_old-change > 1 or < -1) in ARID1A KO hpiNs overlap the previously identified FOS/BAF binding sites (Supplementary Table 4). Of these, the majority (n = 1393, 76%) display decreased chromatin accessibility in the absence of ARID1A (Fig. 3c). ARID1A and ARID1B are mutually exclusive subunits required for the assembly of the canonical BAF complex; they connect the core subunits to the ATPase module^46^, and recently they have been shown to be critical for BAF-protein network interactions^27^. Despite playing a similar role in the canonical BAF complex, they are not interchangeable, as variants in *ARID1A* or *ARID1B* cause a severe neurodevelopmental phenotype^2^. Thus, the decrease in chromatin accessibility at a subset of FOS binding sites in ARID1A KO hpiNs suggests that either there is insufficient canonical BAF complex available, or ARID1B-containing BAF complexes are unable to fully compensate for ARID1A KO at all FOS/BAF binding sites.

BAF is known to interact with TFs in addition to FOS^29^. Therefore, when ARID1A function is disrupted, we were not surprised to identify non-FOS-bound regions across the neuronal genome that also display differential chromatin accessibility. Across the entire genome, including FOS-bound and non-FOS-bound sites, there were more regions with decreased chromatin accessibility versus increased chromatin accessibility in the ARID1A KO hpiNs (Extended Data Fig. 5c). The log_2 f_old-change of chromatin accessibility relative to wildtype was correlated between the ARID1A KO and het hpiNs at all time points (Fig. 3d), suggesting that the ARID1A KO cells adequately model the heterozygous state seen in human patients. As the chromatin accessibility changes were more pronounced in the ARID1A KO versus the ARID1A het hpiNs, we focused our analysis on the ARID1A KO hpiNs.

Given that BAF has been shown to regulate AP-1 TF expression in some cell types^47,26,48–50^, we considered the possibility that the decreased accessibility at FOS/BAF binding sites in the ARID1A KO hpiNs was driven by decreased expression of FOS. However, we found that ARID1A KO hpiNs have increased *FOS* RNA expression and a trend towards increased FOS protein before and 2 hr after depolarization, as compared to wildtype (Fig. 3e, Extended Data Fig. 5d). Likewise, in the absence of ARID1A, RNA expression of the activity-inducible AP-1 TF *JUNB* is increased after neuronal depolarization. By contrast, RNA expression of the constitutively expressed AP-1 TF *JUN* is decreased at all timepoints (Extended Data Fig. 5d). Thus, while we cannot exclude the possibility that decreased *JUN* expression contributes to the decrease in the accessibility of FOS/BAF binding sites, it is likely that activity-dependent decreases in chromatin accessibility at FOS/BAF binding sites are largely due to the inability of the BAF complex to effectively remodel chromatin after recruitment of FOS.

We next asked whether the decrease in chromatin accessibility at FOS/BAF binding sites in ARID1A KO hpiNs affects the expression of nearby genes. We found that genes near FOS/BAF-bound sites with decreased accessibility are more likely to have decreased expression in the ARID1A KO hpiNs (Fisher’s exact test, -KCl and 2 hr p < 2.2e-16), suggesting a direct effect of decreased chromatin accessibility on nearby gene expression. Overall, there are 340 genes with decreased expression that are near at least one FOS/BAF-bound region with decreased accessibility post-membrane depolarization in ARID1A KO hpiNs (Supplementary Table 4). Of these, 107 (31%) are activity-induced genes, and 33 (10%) are ASD-associated genes, including 11 genes that are both activity-induced and ASD-associated (*CD276, CELF4, CPEB4*, *DST, EPH2B, MAPK8IP1, PCDH9, SACS, SDC2, TBL1XR1, YWHAZ*). *SACS* and *SDC2* are two example genes with activity-induced expression in wildtype but not in ARID1A KO hpiNs (Fig. 3f-g). *SACS* encodes the Sacsin protein, which acts as a protein chaperone with diverse functions^51^. Biallelic variants in SACS cause autosomal recessive spastic ataxia of Charlevoix-Saguenay (ARSACS)^52^, and *de novo* heterozygous loss-of-function and missense variants in *SACS* have been identified in individuals with ASD ^53–55^, leading to classification as a strong candidate ASD gene. *SDC2*, another strong candidate ASD gene, encodes a transmembrane heparan sulfate proteoglycan important for dendritic spine formation and maturation^56,57^, and rare heterozygous variants in *SDC2* have been associated with ASD^58,59^. Thus, the disruption of ARID1A function leads to dysregulation of a subset of activity-regulated genes, with effects on neurodevelopment as well as neuronal function throughout life.

### Rare variants within FOS/BAF binding sites in individuals with ASD can disrupt enhancer function

Given the clear importance of the FOS/BAF-mediated activity-dependent gene program for normal neurodevelopment, we hypothesized that FOS/BAF binding sites would be highly conserved across mammalian evolution. However, using phyloP scores, a measure of evolutionary conservation, we found that the 500 bp FOS/BAF binding sites are largely under neutral evolution (Fig. 4a). Since sequence alterations in the canonical AP-1 motif strongly influence FOS binding and enhancer activation^60^, we next considered the possibility that the seven bp AP-1 motifs within the FOS/BAF-bound regions might be more conserved than the larger 500 bp FOS/BAF binding site. Indeed, we found that many of these AP-1 motifs are well-conserved, illustrating the importance of the AP-1 motif within the larger FOS/BAF-bound region and indicating a likely critical function for these sites in neurodevelopment across many species (Fig. 4a).

**Figure 4.**
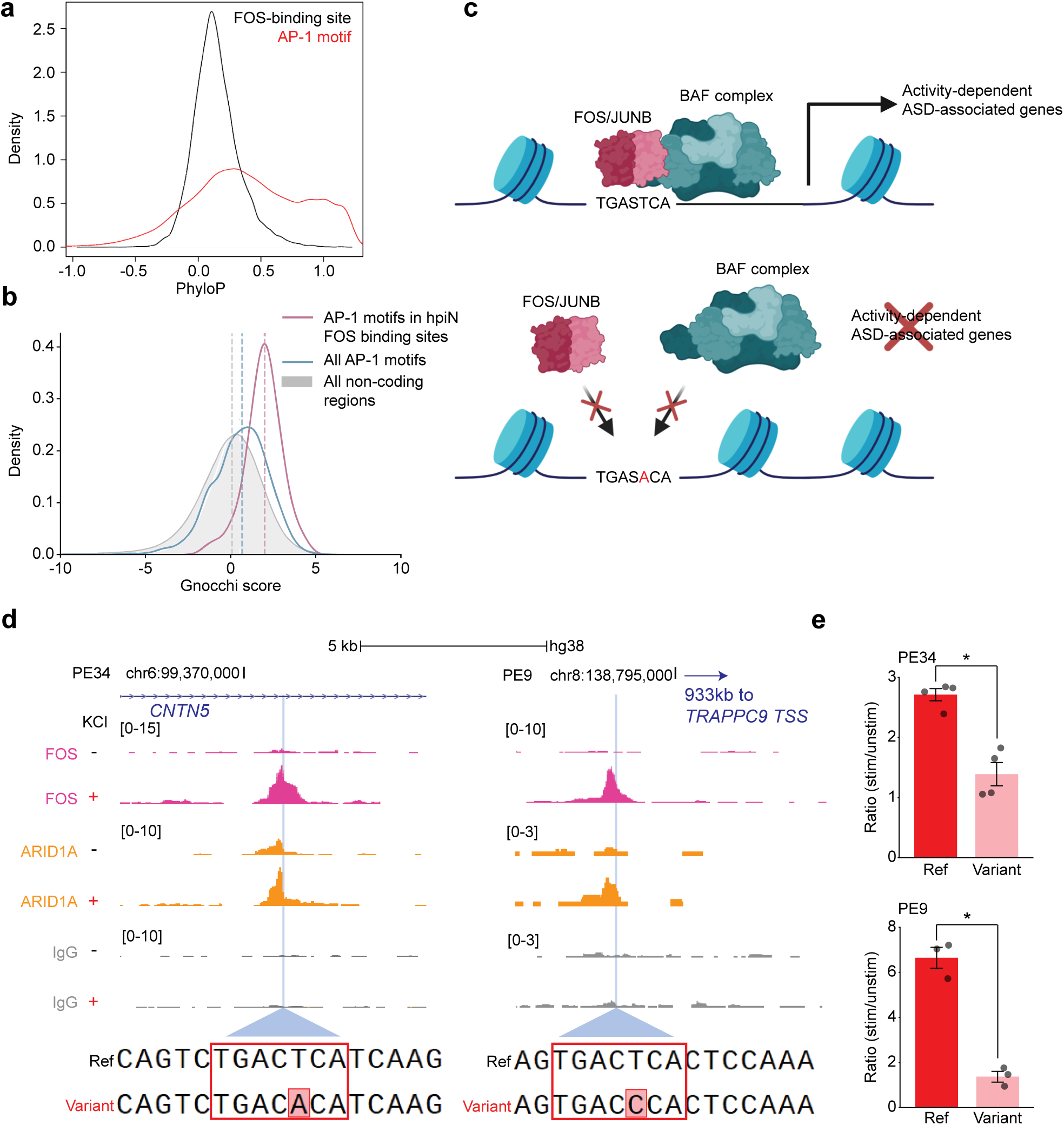
SNVs in the AP-1 motif can disrupt activity-dependent enhancer function. (**a**) Distribution of the phyloP scores for the 500 bp FOS/BAF binding sites (black) and the AP-1 motifs within FOS/BAF binding sites (red). (**b**) Distribution of non-coding constraint (Gnocchi score) in AP-1 motifs in hpiN FOS/BAF binding sites (pink), all AP-1 motifs across the genome (blue), and all non-coding regions (gray). (**c**) Schematic illustrating how SNVs in the AP-1 motif might impact gene expression and contribute to NDD pathogenesis. (**d**) Genome browser tracks of PE34, in an intronic region of the ASD-associated gene *CNTN5*, and PE9, in an intergenic region closest to the ASD-associated gene *TRAPPC9*, with CUT&Tag signal for FOS (pink, n = 6), ARID1A (yellow, n = 3), and paired IgG negative control (gray) before (-) or 2 hr after (+) depolarization. The light blue line highlights the location of the AP-1 motif with an SNV in position 5. Scale is indicated in the brackets. (e) Bar graph showing the ratio of depolarized versus unstimulated reporter assay signal driven by 500 bp enhancer regions with the reference sequence (dark red) compared to the same 500 bp regions with a single SNV introduced in the AP-1 motif (light red) for PE34 and PE9 (mean ± s.e.m., dots = mean of 2 technical replicates for each biological replicate, n = 4 biological replicates for PE34, n = 3 biological replicates for PE9, *adjusted p<0.05). Cultured mouse E16.5 cortical neurons were transfected on Day 3-4, silenced on Day 5, and depolarized with addition of KCl on Day 6.

Recent work also suggests that neuronal FOS binding sites are frequent sites of evolution between the chimpanzee and human, suggesting a human-specific function for some of these regions^34^. To determine if the AP-1 motifs within FOS/BAF-bound sites are constrained in the human population, we looked for human variation in the gnomAD database within AP-1 motifs across the genome and in neuronal FOS/BAF-bound sites^61^. Overall, we found that AP-1 motifs across the genome are more constrained than other non-coding regions genome-wide (64^th^ percentile) and that AP-1 motifs within neuronal FOS/BAF binding sites are highly constrained (88^th^ percentile) (Fig. 4b). This high degree of constraint suggests that any changes in the AP-1 motifs in FOS/BAF binding sites are likely to be deleterious and contribute to human disease (Fig. 4c).

To further investigate this possibility, we queried the SFARI Simons Foundation Powering Autism Research for Knowledge (SPARK) dataset, which contains whole-genome sequencing data for 3,189 individuals with ASD and 8,360 non-ASD individual samples in 3,088 families^62^. Within the characterized hpiN FOS/BAF binding sites, we identified 266 rare (max allele frequency < 0.0005) SNVs located at conserved nucleotide positions in AP-1 motifs (Supplementary Table 5). There was no significant enrichment of AP-1 motif variants in ASD versus non-ASD individuals within the cohort, suggesting that further functional analysis was needed to determine which variants might be benign or deleterious.

Towards this end, we selected 10 SNVs for functional characterization, prioritizing variants that are highly conserved across evolution, have high constraint scores in the human population, are near ASD-associated genes or genes known to be important in neurodevelopment, and that are apparently *de novo* in the affected individual or are inherited but absent in unaffected siblings. We introduced these putative enhancers (PE) into a luciferase reporter plasmid along with a minimal promoter, and the effect of the sequence variants on enhancer-driven activity-dependent gene expression was assessed in cultured mouse cortical neurons. Across all the variants tested, there is a significant genotype effect by ANOVA analysis (p = 0.0005), with decreased activity-dependent reporter expression for the allele with the SNV versus its corresponding reference sequence (Extended Data Fig. 6). Two variants (in PE34 and PE9) have a statistically significant effect on activity-dependent reporter expression (Fig. 4d-e, adjusted p<0.05). Strikingly, the five SNVs with the biggest difference in reporter expression between the reference and variant sequences are all in the third or fifth positions of the AP-1 motif, consistent with prior data showing that SNVs in those positions are most likely to disrupt AP-1 binding and eliminate enhancer formation in mouse fibroblasts^60^.

The first variant (in PE34) with a statistically significant impact on activity-dependent enhancer function is in an intronic region near the ASD-associated gene *CNTN5*, which encodes a cell adhesion molecule. Copy number variants (CNVs) affecting *CNTN5* are believed to be an intermediate risk factor for ASD, and intronic deletions in particular are associated with an increased risk for ASD in families^63^. The AP-1 variant identified in the SPARK dataset is a heterozygous substitution of A for T in the fifth position of the AP-1 motif (chr11:99371063 T>A, gnomAD mAF = 0). This variant was apparently *de novo* in an individual with non-verbal ASD, last evaluated at five years of age, with no self-reported or clinically causative coding region variant identified. Interestingly, the AP-1 variant overlaps with 26 of 30 regions of large deletions or duplications (average size 18 Mb) involving the *CNTN5* locus that were previously reported from the DECIPHER database. Sixteen of the 26 individuals with these deletions had reported developmental delays, intellectual disability, and/or ASD^63^. These results support the hypothesis that the presence of this AP-1 variant results in the mis-regulation of *CNTN5* expression, which could increase the risk for or be a cause of ASD and other NDDs.

The second variant with a statistically significant impact (in PE9) is in an enhancer within an intergenic region nearest to the ASD-associated gene *TRAPPC9*, which encodes a protein that has been shown to be important for NF-kappa-B signaling^64^. Variants in *TRAPPC9* are associated with autosomal recessive intellectual disability, at times with additional features including facial dysmorphisms, obesity, and hypotonia^65^. The enhancer variant identified in the SPARK dataset is a heterozygous substitution of C for T in the fifth position of the AP-1 motif (chr8:138793494 T>C, gnomAD mAF = 2e-4). This AP-1 variant was detected in an individual with ASD, who inherited the variant from an unaffected mother. This patient was last evaluated at nine years of age, with no self-reported or clinically causative coding region variant identified. A study in mice suggests that there is parent-of-origin bias in the expression of *TRAPPC9*, such that 70% of expression is from the maternally inherited allele^66^. However, others have found that the degree of imprinting may be cell-type-specific and variable^67^. Nevertheless, due to imprinting, a heterozygous AP-1 variant in the maternally inherited allele could have a more significant impact on gene expression in the child, resulting in a substantial decrease in *TRAPPC9* expression, increasing risk for neurodevelopmental abnormalities.

Taken together, our findings suggest that rare non-coding variants in AP-1 motifs in FOS/BAF-regulated regions can disrupt enhancer function in human neurons and are a candidate contributing factor in ASD pathogenesis.

## Discussion

The BAF complex is important for normal neurodevelopment, and mutations in many BAF subunits lead to neurodevelopmental disorders, including ASD^2^. Yet the factors that direct BAF binding in human post-mitotic neurons were previously not known. Here we demonstrate that BAF is critical to the activity-dependent regulation of FOS binding sites and that perturbation of the activity-dependent gene program likely contributes to the pathogenesis of ARID1A- and other BAF subunit-related disorders. Additionally, we find that rare ASD patient SNVs in the AP-1 motif in FOS/BAF-bound regions disrupt activity-dependent enhancer function, with the potential to affect expression of nearby ASD-associated genes. Further research into the role and impact of SNVs in AP-1 motifs within neuronal BAF-regulated enhancer regions could improve our understanding of modifying genetic risk factors for ASD.

A large knowledge gap exists regarding the role of rare variants in the noncoding genome in ASD pathogenesis, in part due to the difficulty in predicting the impact of any given variant on enhancer function^11^. Prior studies, including those from our group, found an overall heritability enrichment for neuropsychiatric disease in neuronal activity-regulated regions^12,13^. The present study builds a mechanistic framework for this observation and suggests that particular attention should be paid to variants in AP-1 motifs within neuronal BAF-bound enhancer regions. In conjunction with computational and machine learning approaches in human genomics, we argue that careful dissection of the functional effects of AP-1 motif variants in the developing brain will shed light on the role of noncoding variation as a driving or modifying genetic risk factor for NDD. More broadly, we hypothesize that AP-1 motif variants are likely to contribute to non-pathological variation in human cognitive and emotional functioning, and it is clear that the field is poised to address these types of questions using whole genome sequencing and phenotypic data from large-scale projects like the UK Biobank and All of Us.

Our findings also add to growing evidence for a model where, in response to neuronal activity, FOS recruits BAF to neuronal enhancer regions to facilitate enhancer formation, chromatin opening, and the expression of downstream genes^4,7^. Furthermore, the changes in FOS/BAF binding sites and nearby genes in ARID1A KO hpiNs suggest that aberrant activity- and FOS-dependent enhancer activation contributes to the pathogenesis of ARID1A-related disease, and perhaps other BAF-related disorders. This has important implications as we consider molecularly targeted therapies in this population. A critical question is to what extent the neurodevelopmental phenotype will be reversible. Our work indicates that in individuals with ARID1A haploinsufficiency there will likely be some fundamental differences in FOS-dependent enhancer regulation throughout early development, which may not be reversible at a later age. However, we also find a role for ARID1A in ongoing activity-dependent processes, which implies that there is potential to augment certain activity-dependent functions, like learning and memory, throughout life. Further work will be required to better understand the reversibility of ARID1A-related ASD phenotypes, and it will be important to extend this work to other BAF complex subunits implicated in human neurological disorders.

More broadly, our results fit into a larger body of work that suggests that perturbation of neuronal activity-dependent gene transcription is an important contributor to ASD pathogenesis. Components of the signaling pathways upstream of FOS have previously been implicated in syndromic and non-syndromic ASD. Examples include the L-type voltage-gated calcium channel (*CACNA1C*, Timothy syndrome)^4^, the calcium/calmodulin-dependent protein kinase CaMKII (*CAMK2A*, *CAMK2G*)^68,69^, the RSK2 serine/threonine kinase in the RAS/MAPK signaling pathway (*RPS6KA3*, Coffin-Lowry syndrome)^70^, and CREB-binding protein (*CREBBP*, Rubinstein-Taybi syndrome)^71^. Our work builds upon this framework by directly implicating mis-regulation of FOS function and downstream gene programs in BAF-related disorders. Along with other work suggesting that BAF complex composition and activity are affected by neuronal activity^49,72^ and that BAF complex perturbation can affect IEG expression^48,50^, our findings emphasize the importance of understanding the complex interplay between neuronal activity, canonical IEGs like FOS, and the BAF complex in the titration of gene expression during normal and abnormal neurodevelopment.

Finally, an important feature of this study was our ability to work directly with human neurons. The non-coding genome is more evolutionarily divergent than the coding regions and therefore less tractable to study in animal models^73,74^. Large-scale sequencing of human postmortem and surgical brain tissue has advanced our understanding of transcriptional regulation across development^75,76^, but neuronal activity-dependent processes, which occur over minutes to hours and vary across cell types, are difficult to study due to the lack of control over neuronal activity in these samples. Here we took advantage of the widely adopted protocol combining *NGN2* overexpression and small molecule patterning to produce upper cortical-like neurons that approximately model upper layer cortical neurons implicated in ASD pathogenesis^77^. This *in vitro* system allows tight control over neuronal depolarization in a relatively homogeneous cell population, as confirmed by our scRNA-sequencing experiments. We and others have used similar differentiation protocols to characterize human activity-dependent neuronal processes^12,13^, and we significantly expand upon this work by exploring the role of the BAF complex in activity-dependent transcription and testing the impact of specific human SNVs on enhancer function.

## Acknowledgements

We are grateful to all the members of the M.E.G laboratory, past and present, for their advice and guidance. We would like to thank the Neurobiology Imaging Facility (NIF) at Harvard Medical School for imaging support and the iPS core at Harvard University for stem cell support. This work was supported by a grant from the Simons Foundation or the Simons Foundation International (703333, M.E.G.). Additionally, we are grateful to all the families in SPARK, the SPARK clinical sites and SPARK staff, and we appreciate obtaining access to phenotypic and genetic data on SFARI Base. S.K.T. received support from NINDS/NIH grants R25NS070682, K08NS130150, and L40NS134078. A.C.C. received support from the Hanna H. Gray Fellowship from the Howard Hughes Medical Institute. C.P.D. received support from NIH fellowships T32-NS007473 and F32-NS112455 and the Harvard Mahoney Neuroscience Institute. J.W.H. received support from the NIH grant NS083524. M.E.G. and R.N.D. are supported by the Allen Discovery Center program, a Paul G. Allen Frontiers Group advised program of the Paul G. Allen Family Foundation. M.E.G. is also supported by the Hock E. Tan and K. Lisa Yang Center for Autism Research at Harvard University.

The content is solely the responsibility of the authors and does not necessarily represent the official views of the National Institutes of Health.

Figure 1a: Created in BioRender. Trowbridge, S. (2025) https://BioRender.com/xn4nny8

Figure 5c: Created in BioRender. Trowbridge, S. (2025) https://BioRender.com/4c9scwx

## Author contributions

SKT, GC, ACC, ECG, and MEG conceived of the experiments. SKT, GC, ACC, JEP, JER, GTK, and CPD conducted the experiments. SKT, ACC, SC, RND, DAH, and KJK performed the analyses. JWH provided resources and technical advice. SKT, GC, and MEG wrote the manuscript with input from all the authors.

## Data availability

The sequencing data described in this study will be made available via the NCBI Gene Expression Omnibus (GEO): GSE297424 (ATAC-seq); GSE297425 (CUT&Tag); GSE297716 (RNA-seq); GSE297426 (scRNA-seq).

Publicly available datasets: SFARI ASD-associated genes 01-16-2024 release (https://gene.sfari.org/database/human-gene/); gnomAD v2.1, v3.1.2 (https://gnomad.broadinstitute.org/)

Approved researchers can obtain the SPARK genome sequencing and phenotyping datasets described in this study by applying at https://base.sfari.org.

## Methods

### hESC culture

The protocol for using hESCs was approved by the Harvard Embryonic Stem Cell Research Oversight (ESCRO) Committee, and experiments were compliant with their policies and guidance. The H9 line with a stably introduced inducible NGN2 cassette was previously described^19^. The HUES64 parental line was a kind gift from Daheron L. (Harvard Stem Cell Institute iPS core), and the H1 (also known as WA01) parental line was purchased from WiCell. All hESCs exhibited normal karyotype, as assessed by both hESC Genetic Analysis Kit (STEMCELL) and Karyostat Assays (Thermo Fisher). Mycoplasma was undetectable with the LookOut Mycoplasma PCR Detection Kit (Sigma).

Tissue culture-treated plates were coated with hESC-qualified matrigel (Corning) diluted in cold DMEM/F-12 medium (Gibco), according to the dilution factor determined for each lot. Coated plates were warmed for at least 1 hr at 37°C and washed 2x with warm DMEM/F-12 prior to use. Cells were maintained in culture and passaged every 3-4 days with 1 mg/mL Dispase II (Life Technologies) or Gentle Cell Dissociation Reagent (STEMCELL).

### hESC NGN2 differentiation to hpiN

Differentiation of hESCs into hpiNs was described previously^78^. At day 0, cells were treated with Accutase (StemPro Accutase, Life Technologies) and plated as single cells at 50,000 cells/cm in mTeSR Plus media (STEMCELL Technologies #05825) supplemented with 10 µM Y-27632 (STEMCELL Technologies) on tissue culture plates coated with 336.67 µg/mL Growth Factor Reduced (GFR) matrigel (Corning). On day 1, the medium was replaced with KSR media (recipe below) supplemented with 100 nM LDN193189 (LDN, STEMCELL Technologies), 2 µM XAV939 (XAV, STEMCELL Technologies), 10 µM SB431542 hydrate (SB, Sigma), and 2 µg/mL doxycycline (Dox) hyclate (Sigma). Day 2 media was 50% KSR media/50% NIM media (recipe below) supplemented with LDN/XAV/SB/Dox. Day 3 media was NIM media supplemented with 2 µg/mL Dox. At day 4, cells were treated with Accutase and plated as single cells at 40,000 cells/cm^2^ (30,000 cells/cm^2^ for HUES64 NGN2) in NB media (recipe below) supplemented with 1X B27 without Vitamin A (Life Technologies), 2-2.4 µg/mL mouse laminin (Gibco), 1 µM ascorbic acid (Sigma), 2 µM dibutyryl cyclic-AMP (Sigma), 20 ng/mL brain-derived neurotrophic factor (rhBDNF, Peprotech), 10 ng/mL glial-derived neurotrophic factor (rhGDNF, Peprotech), 10 µM Y-27632, and 2 µg/mL Dox on tissue culture plates coated with GFR matrigel. The day 4 media without Y-27632 is referred to as complete NB (cNB) media. On day 5 the media was replaced with cNB media. Thereafter, weekly cNB media changes were performed with additional Dox supplementation in between media changes (days 8, 15 and 22).

#### The following differentiation media were used

KSR media: Knockout DMEM medium (Life Technologies), 15% knockout serum replacement (KOSR, Life Technologies), 2 mM L-Glutamine (Life Technologies), 1X MEM non-essential amino acids (MEM NEAA, Life Technologies), 1X penicillin/streptomycin (pen/strep, Life Technologies) and 1X 2-mercaptoethanol (Gibco); NIM media: DMEM/F-12 medium (Invitrogen), 1X GlutaMAX, 1X MEM NEAA, 1X pen/strep, 0.16% D-glucose (Sigma) and 1X N2 supplement-B (STEMCELL Technologies); NB Base media: Neurobasal medium (without glutamine, Life Technologies), 1X GlutaMAX, 1X MEM NEAA, 1X pen/strep and 1X N2 supplement-B.

### KCl depolarization of differentiated hpiNs

On day 27 post-differentiation, hpiNs were silenced overnight with 1 µM TTX (VWR) and 100 µM AP5 (Tocris) for 16-18 hr. The following day, hpiNs were stimulated with KCl Depolarization Buffer (170 mM KCl, 2 mM CaCl_2,_ 1 mM MgCl_2 a_nd 10 mM HEPES) to the final concentration of 55 mM KCl for 15 min, 1 hr, 2 hr, or 6 hr. A subset of hpiNs were left unstimulated by maintaining them in the silencing media without the KCl treatment. After the stimulation, cells were washed 1x with 1X PBS and collected in the buffers associated with each assay.

### gRNA for genome engineering

The gRNA sequence used to introduce *TRE3G-NGN2* (5’-GGGGCCATAGGGACAGGAT-3’) was described previously^19^. For other gRNAs, the Online Synthego CRISPR Design Tool was used to design and select guides with high on-target scores and few potential off-target sites. The ARID1A gRNAs targeted the second exon of *ARID1A* (guide 3 = 5’-GTCCAGTCCAATGGATCAGA-3’, guide 4 = 5’-GTAGTCCCGCCATATGGCTG-3’). The FOS gRNA targeted the second exon of *FOS* (guide 3 = 5’-GGTCTGCGATGGGGCCACGG-3’). TRE3G-NGN2 gRNA was synthesized by Synthego. The template oligos for other gRNA’s above were ordered from IDT, and gRNAs were synthesized with the MEGAScript T7 Transcription Kit (Invitrogen) according to the manufacturer’s instructions.

### Engineering hESCs (FKO, ARID1Bhet, HUES64/H1 NGN2 lines)

Stem cells at 70-80% confluency were enzymatically dissociated into single cells with Accutase (Thermo Fisher, A1110501) for 15-20 min at 37°C. In parallel, RNP complex was prepared by mixing and incubating Cas9 protein (1 µg/µL, Pna Bio, CP01), gRNA (2-3 µg/µL) and donor plasmid (3-4 µg/µL) per 1x10^6^ cells for 10 min at room temperature. RNP complexes for generating ARID1A mutant and FOS KO lines did not contain any donor plasmid. Dissociated cells were resuspended in the Resuspension Buffer R from the Neon Transfection System 100 µL Kit (Thermo Fisher, MPK10096) and aliquoted to have 2x10^6^ cells for each reaction. RNP complexes were mixed with each cell aliquot, and nucleofection was performed using Neon Transfection System (Thermo Fisher, MPK5000) according to the manufacturer’s instructions. An efficient Neon system protocol for nucleofecting each stem cell line was first optimized using a pmaxGFP vector (Lonza) as a donor plasmid: H9 NGN2=protocol 2, H1=protocol 2, and HUES64=protocol 3. The nucleofected cells were plated on warm hESC-qualified Matrigel-coated 100 mm tissue culture plates with mTeSR Plus media supplemented with Y-27632 or CloneR (STEMCELL).

#### For TRE3G-NGN2

Media was changed daily with fresh mTeSR Plus media until cells were approximately 70% confluent. Cultures were selected with an overnight puromycin (1 µg/mL, STEMCELL, 73342) treatment and maintained in mTeSR Plus media until the size of each colony was about 1/5 of field of view under 10X objective. Each clonal colony was picked into one well of hESC-qualified Matrigel-coated 96-well plate and maintained in mTeSR Plus media. During the next passage into a new 96-well plate, genomic DNA was extracted from each well using a Genomic DNA Purification Kit (STEMCELL, 79020) for genotyping. Genotyping to verify homozygous knock-in of the TRE3G-NGN2 construct was performed with PCR using the primers described previously^19^: 5’ junction (5’-CTCTAACGCTGCCGTCTCTC-3’ and 5’-TGGGCTTGTACTCGGTCATC-3’), 3’ junction (5’-CACACAACATACGAGCCGGA-3’ and 5’-ACCCCGAAGAGTGAGTTTGC-3’) and locus (5’-AACCCCAAAGTACCCCGTCT-3’ and 5’-CCAGGATCAGTGAAACGCAC-3’).

#### For FOS and ARID1A

Media was changed daily with fresh mTeSR Plus media for two days. Cultures were treated with Accutase, and 2000 cells were plated in hESC-qualified Matrigel-coated 100 mm plates containing mTeSR Plus supplemented with CloneR (mTeSR Plus+CloneR). Media changes were performed the next day with mTeSR Plus+CloneR and then with mTeSR Plus for the following days until each colony was about 1/5 of field of view under the 10X objective. Each clonal colony was picked into one well of a hESC-qualified Matrigel-coated 96-well plate and maintained in mTeSR Plus media. During the next passage into a new 96 wells plate, genomic DNA was extracted from each well using a Genomic DNA Purification Kit. PCR was performed using the primers flanking the gRNA-targeted regions: ARID1A_F (5’-CCAGAGGCCATCAAAGCTC-3’) and ARID1A_R (5’-GTCCTTGCTGCGGTCCTG-3’), FOS_F (5’-CTCTCTTACTACCACTCACCCGC-3’) and FOS_R (5’-GCCATCTTATTCCTTTCCCTTCGG-3’). The PCR products were purified with QIAquick PCR Purification Kit (Qiagen, 28106) and used to perform a TOPO cloning reaction with the Zero Blunt TOPO PCR Cloning Kit (Thermo Fisher, 450245) according to the manufacturer’s instructions. Stbl3 chemically competent cells (Life Technologies, C737303) were transformed with the TOPO cloning products and grown on kanamycin LB plates. The resulting colonies were picked and sent for Sanger sequencing. Lines with frameshift variants predicted to lead to protein truncation were selected and normal karyotypes confirmed with the Karyostat Assay (ThermoFisher).

The final lines used for experiments were as follows: FKO 3-6-19 with one allele with a single bp deletion (chr14:75,279,989 del) and one allele with an 11 bp deletion (chr14:75,279,985-75,279,995 del), predicted to lead to early stop codons in exons 2 and 3, respectively; ARID1A KO A3-25 with one allele with a single bp insertion (chr1:26,729,670 ins A) and one allele with a 10 bp deletion (chr1:26,729,660-26,729,669 del), both predicted to lead to early stop codons in exon 2; ARID1A KO A3-44 with one allele with a 14 bp deletion (chr1:26,729,653-26,729,666 del) and one allele with a 2 bp deletion (chr1:26,729,671-26,729,672 del), both predicted to lead to early stop codons in exon 2; ARID1A het A4-16 with one allele with a 2 bp insertion (chr1:26,729,694 CC ins), predicted to lead to an early stop codon in exon 2; ARID1A het A4-43 with one allele with a 2 bp deletion (chr1:26,729,693-26,729,694 del), predicted to lead to an early stop codon in exon 2.

### qPCR

Cells were treated with 5 µM nimodipine (Sigma Aldrich # N149) or 5 mM EGTA (Boston BioProducts # BM-151) for 1 hr or 10 min, respectively, prior to KCl stimulation. Cells were washed 1x with 1X PBS and collected in Trizol (Invitrogen). Total RNA was extracted with Trizol/Chloroform and purified with RNeasy Micro Kit (QIAGEN, 74104) according to the manufacturer’s instruction. During the purification, membranes on the columns were treated with DNaseI to remove potential DNA contamination. The isolated RNA was quantified using NanoDrop spectrophotometer and used to synthesize complementary DNA (cDNA) with SuperScript VILO cDNA Synthesis Kit (Thermo Fisher, 11754050) according to the manufacturer’s instructions. The cDNA was used to perform qPCR assay with SYBR Green reagent (Applied Biosystems), and the Quant Studio 3 Biosystem (Thermo Fisher) was used to amplify and detect the transcripts in each sample. The expression level of *FOS* transcript was normalized by the level of *PGK1* transcripts detected. The following primer pairs were used: FOS_qPCR_F-5’ TGGCGTTGTGAAGACCATGA 3’, FOS_qPCR_R-5’ CTGTCTCCGCTTGGAGTGTA 3’. PGK1_qPCR_F-5’ CCGCTTTCATGTGGAGGAAGAAG 3’, PGK1_qPCR_R-5’ CTCTGTGAGCAGTGCCAAAAGC 3’.

### Immunohistochemistry

Day 28 hpiNs differentiated on GFR Matrigel-coated German glass coverslips (VWR) were washed once with 1X PBS and fixed with 4% paraformaldehyde (v/v) and 4% Sucrose (w/v) in 1X PBS for 15 min at room temperature. Fixed coverslips were washed 5x with 1X PBS and blocked with 5% donkey serum (DS, v/v) and 0.2% Triton X-100 (v/v) in 1X PBS for 1 hr at room temperature. Coverslips were washed 1x with 1X PBS and incubated in primary antibodies that were diluted in antibody buffer (2% DS and 0.1% Triton X-100 in 1X PBS) overnight at 4°C. The primary antibodies used include: anti-FOS (rabbit, 1:2000, Synaptic Systems, 226003), anti-MAP2 (chicken, 1:2500, Lifespan Biosciences, LS-C61805), and anti-BRN2 (rabbit, 1:1000, CST, 12137S). The next day, coverslips were washed 3x with wash buffer (0.02% Triton X-100 in 1X PBS) and incubated in AlexaFluor-conjugated secondary antibodies that were diluted in antibody buffer (1:1000, Life Technologies) for 2 hr in room temperature in the dark. The coverslips were washed 3x with wash buffer and 1x with PBS. Stained samples were mounted on the Superfrost Plus slides (Thermo Fisher) using DAPI Fluoromount G (Fisher Scientific). Immunofluorescent images were acquired using Olympus FV1000 Confocal Microscope (40x oil 1.3NA objective) or the Leica SP8X STED microscope (40x oil objective) in the Neurobiology Imaging Facility at Harvard Medical School. Acquired images were processed and quantified using Fiji software.

### Bulk RNA-seq collection and library preparation

Each experiment had three biological replicates (independent differentiations), unless otherwise indicated. RNA was harvested from neurons on day 28 of differentiation following the stimulation protocol as described above. For the ARID1A het/KO experiments, samples were collected for two ARID1A het lines, two ARID1A KO lines, and one paired wildtype line. Bulk RNA-sequencing libraries were prepared using the SMARTer Stranded Total RNA-Seq Kit v2 - Pico Input Mammalian kit (Takara Bio, 634411), as per the kit instructions. Ten nanograms RNA input of RNA input was used for each library. Libraries were quantified using Qubit and a BioAnalyzer and sequenced on an Illumina NextSeq 500 using paired-end 37 bp reads.

### Bulk RNA-seq analysis

RNA sequencing reads were aligned using HISAT2 v2.1.0 with the parameters –rna-strandedness RF and –mp 4,2. Reads that mapped were then assigned to features in the Gencode V32 human transcript reference using FeatureCounts (Rsubread v2.0.0)^79^. Differential analysis was then carried out at the gene level using DESeq2 v1.46.0^80^ with default parameters. Differential genes were called as those with an adjusted p-value < 0.05 and FPKM ≥ 2 in at least one sample. Genes on the X and Y chromosomes were filtered out for analyses that compared between male and female lines.

### Neuronal dissociation for 10x single cell RNA-sequencing

After depolarization (-KCl, 2 hr KCl, 6 hr KCl), neurons were dissociated to single cells using the Worthington Papain Dissociation System (Worthington Biochem, LK003150). Neurons were washed twice on tissue culture plates with PBS + 0.5% BSA. Neurons were then incubated for 15 min in Papain + DNAse solution (1 vial Papain + 1 vial DNAse in 10 mL Dissociation Media [HBSS + 10 µM HEPES + 1.72 mg Kynurenic acid + 8.6 mg MgCl2·6H2O +63 mg D-Glucose]+ 5 inhibitors of transcription and translation in order to preserve the stimulated transcriptional state (1 µM Tetrodotoxin citrate, 100 µM D-AP5, 5 µg/mL Actinomycin D, 10 µM Triptolide, 10 µg/mL Anisomycin)^81^. Papain solution was removed and replaced with Dissociation Media containing 0.05% BSA and 1% Ovomucoid powder to inhibit Papain. Cells were then collected in Dissociation Media + 0.05% BSA and resuspended to make a single cell solution. Cells were pelleted at 300xg for 5 min at 4°C and resuspended in Neurobasal media containing the 5 inhibitors and then passed through a 40 µM filter.

### scRNA-seq using 10x genomics

Single neurons dissociated as described above were processed for 10x single cell RNA-sequencing exactly according to the 10x Genomics Chromium Next GEM Single Cell 3’ Reagent Kits v3 (Dual Index) protocol. Briefly, for each timepoint, 16,500 cells were input at a density of 1000 cells/µL for a targeted recovery of 10,000 cells. GEMs were created on a Chromium Controller, followed by library creation exactly as described in the 10x Genomics protocol. Library quality was verified on a Bioanalyzer, and libraries were quantified using the KAPA library quantification kit.

### scRNA-seq analysis

10x libraries were pooled at equal depth and sequenced a total of 3 times on a NextSeq 500 to a final mean read depth of 24,742 (-KCl), 22,647 (2 hr KCl), and 32,405 (6 hr KCl). Reads were processed using CellRanger V6.1.2 mkfastq and cellranger count. Data was read into Seurat V4.1.0, and analyzed and visualized using this package. Based on QC metrics, we filtered for cells with unique features counts greater than 200 and less than 2500, percent mitochondrial reads less than 8% and percent ribosomal RNA reads less than 20%. We then normalized the data using the function NormalizeData(), and scaled the data using ScaleData(). To integrate the three timepoints into one, we used the function IntegrateData().

### CUT&Tag

Protocol was adapted from a previously described protocol^82^. Cells were washed with cold 1X PBS and collected in NE1 Buffer (20 mM HEPES pH 7.9, 10 mM KCl, 0.1% Triton X-100, 3 mM MgCl_2)_ with the following supplements: 0.5mM Spermidine (Sigma), 1X Roche complete Protease Inhibitor EDTA-Free (Sigma) and 10mM Sodium Butyrate (Millipore). Tubes were rotated for 10 min and centrifuged at 4°C for 10 min each to isolate and pellet the nuclei. Nuclei were resuspended in a total of 1 mL WB150 Buffer (20 mM HEPES pH 7.5, 150 mM NaCl, 0.2% Tween-20, 0.1% BSA (w/v)) with supplements, counted, and aliquoted to have 500k nuclei/reaction or 1 million nuclei/reaction (for NPAS4 only). Nuclei were bound on magnetic ConA beads (Bangs Laboratories) that were washed in Binding Buffer (20 mM HEPES pH 7.9, 10 mM KCl, 1 mM CaCl_2,_ 1 mM MnCl_2)_. The ConA bead-bound nuclei were resuspended in Antibody Buffer and incubated with primary antibody at room temperature for 1.5 hr or at 4°C overnight. The following primary antibodies were used at 1:50 dilution: H3K27ac (Abcam, ab4729), FOS (Synaptic Systems 226003), NPAS4^83^, ARID1B (E9J4T, CST 92964S), ARID1A (D2A8U, CST 12354S), SMARCB1 (D8M1X, CST 91735S), SMARCC1 (D7F8S, CST 11956S), and control polyclonal rabbit IgG (CST 2729S). Next, nuclei were resuspended in Antibody Buffer (2 mM EDTA pH 8.0, 0.1% Triton X-100 in WB150 Buffer with supplements) and incubated with the anti-rabbit guinea pig secondary antibody (ThermoFisher, NBP172763) at room temperature for 1 hr.

After antibody incubations, ConA bead-bound nuclei were washed 3x with WB150 Buffer with supplements, resuspended in WB300 Buffer with supplements, and incubated with pA-Tn5 (homemade in lab) at a final concentration of 25 nM at room temperature for 1 hr. Next, nuclei were washed 3x with WB300 Buffer. (20 mM HEPES pH 7.5, 300 mM NaCl, 0.2% Tween-20, 0.1% BSA (w/v)) with supplements, resuspended in Tagmentation Buffer (10 mM MgCl_2 i_n WB300 Buffer with supplements) and incubated for 1 hr for tagmentation. The tagmentation reactions were stopped by directly adding 16.7 mM EDTA pH 8.0, 0.1% SDS and 166.7 µg/mL proteinase K (QIAGEN) and incubating at 50°C for 1 hr. DNA was purified with standard phenol/chloroform/isoamyl ethanol extraction with ethanol precipitation. The purified DNA was resuspended in 20 µL 10 mM Tris pH 8.0.

CUT&Tag libraries were prepared using KAPA HiFi PCR Kit (Roche), closely following the procedure described previously^23^, with the following PCR cycling conditions: 72°C for 5 min, 98°C for 30 seconds, 14 total cycles of 98°C for 10 seconds, 63°C for 30 seconds, and 72°C for 1 minute, followed by 72°C for 5 min. PCR-amplified libraries were purified in three steps by adding 1) 0.5X ratio of AmPureXP beads, followed by collection of the supernatant containing unbound small fragments; 2) 1.3X ratio of AmPureXP beads, two washes in 80% ethanol, and elution in 53 µL 10 mM Tris pH 8.0; and 3) 1.1X ratio of AmPureXP beads, two washes in 80% ethanol, and elution in 20 µL 10 mM Tris pH 8.0. Libraries were quantified and characterized with KAPA Library Quantification Kit (Roche). Final library products were pooled and sequenced in a NextSeq500 (Illumina) using paired-end 37 bp reads.

### CUT&Tag analysis

Sequencing reads were demultiplexed and then trimmed using Trimmomatic v0.36 and kseq. Trimmed reads were aligned to the hg38 genome using Bowtie2 v2.2.9 with the following parameters: -local --very-sensitive-local --no-unal --dovetail --no-mixed --no-discordant --phred33 -I 10 -X 700. Duplicate reads were removed with Picard, and multi-mapped reads were removed with samtools v1.3.1 using the parameter -q 10. Sequencing reads were calibrated for sequencing depth, normalizing to reads per million mapped reads using a script adapted from Skene and Henikoff^84^. To view normalized coverage tracks, merged bigwig files from all relevant biological replicates were uploaded to the UCSC Genome browser. For FOS and NPAS4, peaks were called using MACS2 v2.1.1.20160309 with the following parameters: -f BAMPE -g hs --nomodel --shift -100 --extsize 200. A paired sample with polyclonal rabbit IgG was used as the negative control sample for peak calling. Peaks were merged using bedtools v2.27.1 merge, and then reproducible peaks were defined by taking those peaks that were present in at least three biological replicates (three independent differentiations). Reproducible peaks that had significantly increased or decreased number of reads at 2 hr post-membrane depolarization, as compared to the unstimulated condition, were determined by using DESeq2 v1.30.1^80^, with an adjusted p-value cut-off < 0.05. Given that FOS and NPAS4 have very low levels of expression in the unstimulated condition, adjusted DESeq2 size factors were calculated using sequencing read depth. Peaks were then centered on the 50 bp region with the maximum CUT&Tag signal in the peak, and 225 bp of flanking sequence were added on both sides to generate uniform 500 bp peaks. For FOS peaks, the presence and location of canonical AP-1 motifs was determined with Homer v4.10.3 annotatePeaks.pl with a custom motif file for the sequence “TGASTCA”, with no mismatches allowed. Other motif analyses were performed with Homer v4.10.3^85^ findMotifsGenome.pl and FIMO v5.5.7^86^. Distance to the nearest TSS and genomic annotations were determined using Homer v4.10.3^85^ annotatePeaks.pl. CUT&Tag signal over regions of interest was determined using Homer v4.10.3 annotatePeaks.pl with merged bam files from the relevant biological replicates. Target CUT&Tag signal was normalized to matched polyclonal rabbit IgG CUT&Tag signal by dividing target signal by IgG signal, if IgG signal > 1. If IgG signal < 1, the raw target signal was used. The ratio of the stimulated to unstimulated CUT&Tag signal was calculated in a similar fashion, by dividing the stimulated signal by the unstimulated signal if the unstimulated signal > 1; if the unstimulated signal < 1, the raw stimulated signal was used. CUT&Tag data from the FOS KO hpiNs was compared to paired wildtype CUT&Tag data collected in the same experiments, to control for variability between hpiN differentiations. CUT&Tag signal over selected regions was visualized with DeepTools v3.5.0^87^ using Reads Per Genome Coverage (RPGC) normalization.

### ATAC-seq

ATAC-seq was performed using a modified protocol as previously described^88^. Neurons were stimulated with KCl on day 28 for the indicated amount of time using the protocol described above. Neurons were washed twice with PBS and then 500 µL Lysis Buffer (10mM Tris-HCl pH 7.4, 10mM NaCl, 3mM MgCl_2,_ 0.1% NP-40, 0.1% Tween) was added to cells on the plate. Cells were scraped into 1.5 mL Eppendorf tubes and rotated for 5 min at 4°C. Nuclei were then pelleted for 10 min at 830xg at 4°C. Supernatant was removed, and pellet was resuspended in 1 mL Lysis Buffer and nuclei were counted. 50,000 nuclei per sample were aliquoted in a new tube and centrifuged 10 min at 830xg at 4°C. Nuclei were then resuspended in transposition mix consisting of 25uL 2x TD buffer (20 mM Tris-HCl pH 7.6, 10 mM MgCl_2,_ 20% Dimethyl Formamide), 0.4 uL Tn5 (100 nM final), 16.5 µL 1x PBS, 0.5 µL 1% Digitonin, 0.5 µL 10% Tween-20, and 5 µL H2O and incubated at 37°C for 30 min on a thermomixer. Following transposition, reaction was cleaned up using the Zymo DNA Clean and Concentrator Kit (Zymo Research D4004) and eluted in 20 µL.

For library preparation, 20 µL eluted DNA was mixed with 25 µL NEB Next 2x PCR Master Mix (New England BioLabs M0541S), and 2.5 µL Ad1 and Ad2 (adapter sequences listed in Buenrostro *et al*., *Nature* 2015^89^). PCR reactions were run with the following conditions. 72°C for 5 min, 98°C for 30 seconds, 8 cycles of 98°C for 10 seconds, 63°C for 30 seconds and 72°C for 1 min. PCR reactions were cleaned up using the Zymo DNA Clean and Concentrator Kit. For the ARID1A het/KO and paired wildtype samples, a double-sided bead purification was used for reaction clean-up instead of the Zymo kit. Briefly, 0.5x volume of AMPure XP beads were added and the supernatant with unbound small fragments was collected, followed by addition of 1.3x (of the original volume) AMPure XP beads, with two 80% ethanol washes, and elution in 20 µL water. Libraries were quantified using the KAPA Library Quantification Kit (KAPA Biosystems KR0405).

There were three biological replicates (independent differentiations) for each experiment. For the ARID1A het/KO experiments, samples were collected for two ARID1A het lines, one ARID1A KO line, and one paired wildtype line.

### ATAC-seq analysis

For ATAC-seq analysis, H9-derived hpiN libraries were sequenced on a NextSeq 500 and H9-derived hpiN ARID1A het and KO with paired wildtype libraries were sequenced on a NovaSeq 6000 and a NovaSeq X plus to a depth of 10-22 million reads per replicate after duplicate removal. Adapters were trimmed from reads using CutAdapt (v2.5). Reads were then mapped to the hg38 reference genome using Bowtie2 (v2.3.4.3) with the following parameters: -p 8 -S -n 2 -e 70 -m 1 -k 1 -l 70. Reads were de-duplicated and then peaks were called using MACS2 with the following parameters: --nomodel --extsize 200. Only peaks that were called in all three replicates were retained and merged to create the final peak list. Bedtools multicov was used to count reads for each sample in the final peak list. After filtering out regions with low reads (maximum count < 5), DESeq2 v1.30.1^80^ was used to call differential peaks between -KCl timepoint and other timepoints or between genotypes using the default normalization (adjusted p-value < 0.05). ATAC signal over selected regions was visualized with DeepTools v3.5.0^87^ using Reads Per Genome Coverage (RPGC) normalization.

### Western blot

Samples were collected in RIPA buffer (Sigma Aldrich) supplemented with EDTA-free protease inhibitor cocktails (Roche). Samples were quantified with the Bio-Rad Protein Assay, and 20 µg protein was boiled in LDS sample buffer (Life Technologies) supplemented with 5% 2-mercaptoethanol (Sigma) for 5 min at 95°C. Prepared protein lysates were separated across 4-12% NuPAGE Bis-Tris Gel (for FOS, Life Technologies) or 3-8% Tris-Acetate Gel (for ARID1A, ThermoFisher) and transferred to 0.45 µm nitrocellulose membranes. Membranes were in blocking buffer (TBST with 5% milk) for 1 hr at room temperature before incubating in primary antibody overnight at 4°C. Primary antibodies used were anti-FOS (rabbit, 1:1000, Synaptic Systems 226008), anti-ARID1A (rabbit, 1:1000, D2A8U CST 12354S), anti-GAPDH (rabbit, 1:5000, Millipore G9545), and anti-H3 (rabbit, 1:2000, Abcam ab1791). Membranes were washed and then incubated in secondary antibody conjugated with horseradish peroxidase (1:10,000, CST 7074S) before being developed or analyzed on BioRad Universal Hood II. Images were analyzed using Fiji software.

### Human constraint analysis

Non-coding constraint analysis was performed as previously described^61^, with scores calculated over 1 kbp regions, created by combining 7 bp AP-1 motifs. 339,053/462,326 (73.3%) of all AP-1 motifs across the human genome were scored. 8,343/11,197 (74.5%) of AP-1 motifs in hpiN FOS-bound regions were scored.

### Variant analysis

Genomic variants in individuals with ASD were identified from the SPARK cohort of 11,545 individuals in 3,087 families (Release Nov. 2021; see SPARKForAutism.org). Variants were filtered using the previously described pipeline^90^. Briefly, variants were filtered based on the following quality metrics (PASS filter, and GQ > 20), maximum population allele frequency in gnomAD v2.1 of 0.0005, and a maximum cohort allele frequency in the SPARK dataset of 0.005. To enrich for variants with the greatest likelihood of impacting functionality, variants were classified using multiple different conservation-based variant effect predictors (e.g., GERP++, CADD, DANN, FATHMMnc) along with RefSeq and Gencode v28 gene annotations. Additional annotations included data from Epigenomics Roadmap and JASPAR TF motifs. All variants meeting the above QC metrics were filtered for GERP > 2 and (CADD > 15 or DANN > 0.85 or FATHMMnc > 0.85). Rare variants found across multiple families in the cohort were filtered out, given the possibility that these were due to a recurrent sequencing error.

### Luciferase assay

The 500-bp FOS binding sites with the reference sequence or introduction of an SNV were ordered as gene blocks (Twist Bioscience) and were cloned into the pNL3.2[NlucP/minP] vector (Promega) using the NEBuilder HiFi DNA Assembly mastermix (NEB), as per the manufacturer’s instructions.

Experiments were performed in cultured mouse cortical neurons, due to higher efficiency of plasmid transfection than in cultured human neurons. E16.5 C57BL/6 mouse embryonic cultures were prepared and cultured as described previously^17^, where dissected neurons were plated at 200,000 cells/well on poly-D-lysine (Sigma) and laminin (Life Technologies)-coated 24 wells tissue culture plates. Experiments were approved by the Animal Care and Use Committee at Harvard Medical School. Mouse cortical cultures were maintained in mNB medium (1X penicillin/streptomycin, 1X GlutaMAX, 1X B27 supplement in Neurobasal medium without glutamine). On days 3 or 4, the pNL3.2[NlucP/minP] vector containing reference or variant sequences were co-transfected with the pGL4.53[luc2/PGK] vector as a transfection control, as well as pBluescript SK(+) and an FUGW-ubi::EGFP vector (Addgene #14883). Transfection was performed with Lipofectamine Stem Transfection Reagent (Thermofisher) according to the manufacturer’s protocol. Neurons were silenced with 1 µM TTX and 100 µM AP5 for 16-18 hr on day 5. On day 6, neurons were depolarized for 6 hr with 55 mM KCl, washed once with 1X PBS(-/-) and lysed with 1X Passive Lysis Buffer (Promega) for at least 45 min while shaking at 225 RPM at room temperature. Luciferase assays were performed on the protein lysates using the reagents from the Nano-Glo Dual-Luciferase Reporter Assay System (Promega), and luminescence from each sample was measured by BioTek synergy 4 microplate reader. Three biological replicates with two technical replicates each were performed for all the constructs. Each biological replicate was performed with paired positive (enh 39)^17^ and negative (pNL3.2 with no enhancer) controls. Luminescence was normalized to the luciferase transfection control and the negative control for all samples. Experiments in which the normalized induction of the positive control < 10-fold were discarded.

