## Supplemental figures for "The BAF complex works with FOS to regulate human neuronal activity-dependent ASD-associated gene programs"

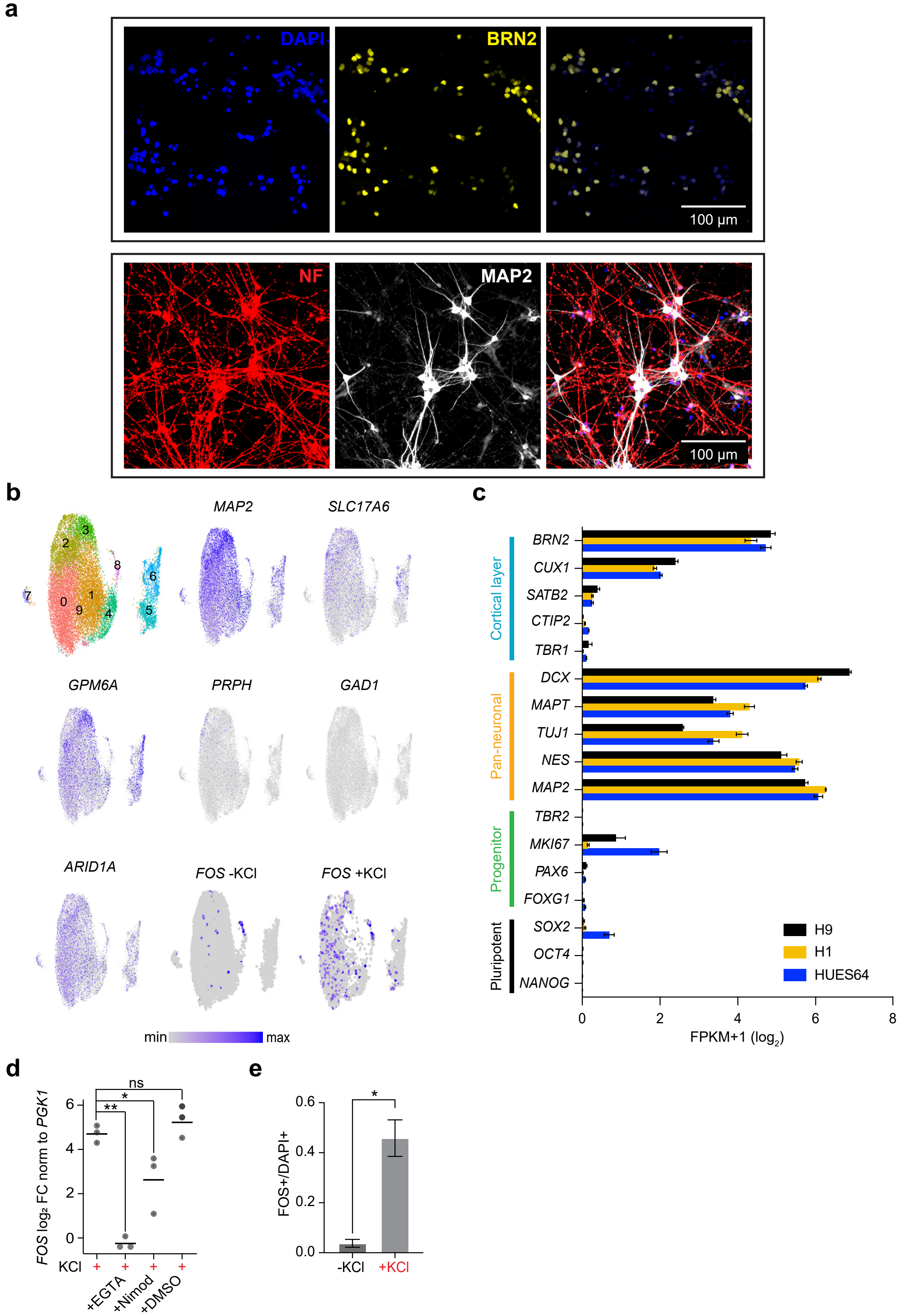


**Extended Data Figure 1.** Characterization of hpiNs. (**a**) Immunohistochemistry for the upper cortical layer marker BRN2 (yellow) and neuronal markers neurofilament (NF, red) and MAP2 (white) in H9-derived hpiNs on Day 28. (**b**) H9-derived hpiN single-cell RNA-seq UMAP plots indicating expression of select genes across the 10 clusters. Expression of *FOS* is shown before (-KCl) and 2 hr after depolarization (+KCl). *MAP2* = neuronal marker; *SLC17A6* = excitatory neuronal marker; *GPM6A* = CNS marker; *PRPH* = PNS marker; *GAD1* = inhibitory neuronal marker; *ARID1A* = BAF complex subunit. (**c**) RNA expression ($\log_2$ (FPKM+1)) of marker genes across hpiNs derived from H9, H1, or HUES64 (n = 3 biological replicates per line, mean ± s.e.m.). (**d**) qPCR quantification of *FOS* mRNA induction in hpiNs 1 hr post-depolarization, after treatment with nimodipine, EGTA, or DMSO control. Normalized signal $\log_2$ fold-change compared to the corresponding unstimulated control in each condition (n = 3 biological replicates; bar = mean of biological replicates, dot = mean of 3 technical replicates for one biological replicate, *p<0.05, **p<0.005, ns = not significant). (**e**) Quantification of FOS+/DAPI+ nuclei from immunohistochemistry in hpiNs before (-KCl) or 2 hr after depolarization (+KCl) (mean ± s.e.m., n = 11 images across 4 biological replicates, *p<0.05, Mann-Whitney test).

**a**

**b**

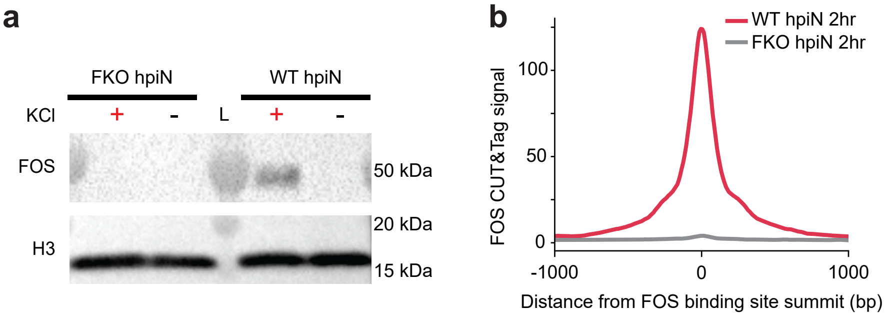


**c**

| | H1 | HUES64 |
|---|---|---|
| Inducible binding sites | 7719 | 1040 |
| Contains canonical AP-1 motif | 6138 (80%) | 923 (89%) |
| Overlaps an H9 FOS binding site | 4553 (75%) | 877 (95%) |

**d**

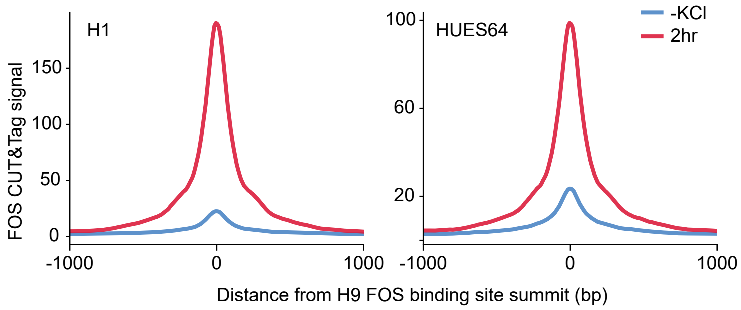


**Extended Data Figure 2.** Validation of activity-dependent FOS binding sites in hpiNs. (**a**) Western blot for FOS from Day 28 wildtype (WT) versus FKO hpiNs, with H3 as the loading control before (-) or 2 hr after (+) KCl depolarization. (**b**) Aggregate enrichment plots of FOS CUT&Tag signal 2 hr after depolarization in WT (red) versus FKO (gray) (n = 3 biological replicates). (**c**) Table showing the number of binding sites in hpiNs derived from H1 or HUES64 hESC and overlap with sites in H9-derived hpiNs. (**d**) Aggregate enrichment plots of FOS CUT&Tag signal from H1 or HUES64-derived hpiNs before (blue) or 2 hr after (red) depolarization over all FOS binding sites from H9-derived hpiNs.

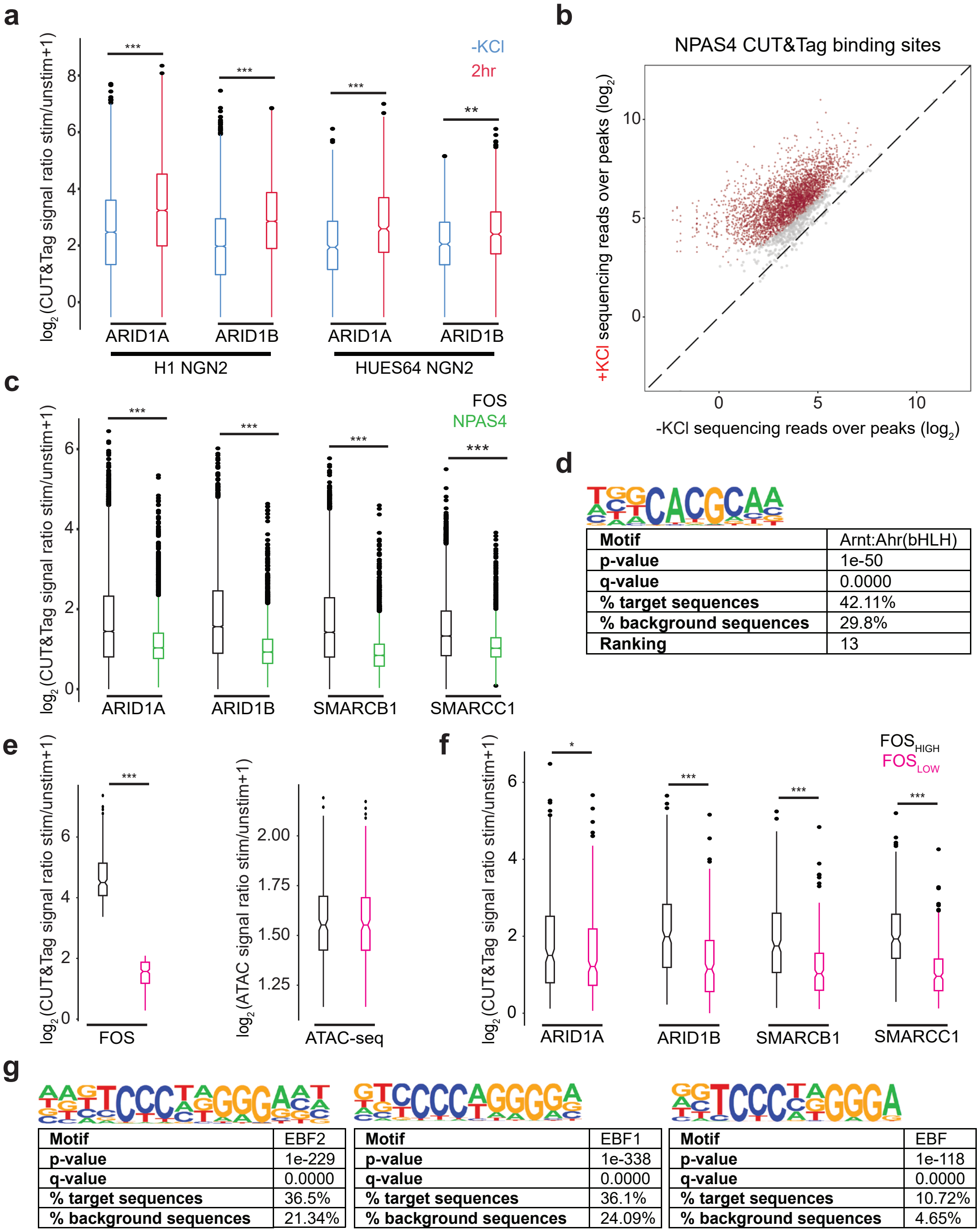


| Motif | Arnt:Ahr(bHLH) |
|---|---|
| p-value | 1e-50 |
| q-value | 0.0000 |
| % target sequences | 42.11% |
| % background sequences | 29.8% |
| Ranking | 13 |

| Motif | EBF2 |
|---|---|
| p-value | 1e-229 |
| q-value | 0.0000 |
| % target sequences | 36.5% |
| % background sequences | 21.34% |

| Motif | EBF1 |
|---|---|
| p-value | 1e-338 |
| q-value | 0.0000 |
| % target sequences | 36.1% |
| % background sequences | 24.09% |

| Motif | EBF |
|---|---|
| p-value | 1e-118 |
| q-value | 0.0000 |
| % target sequences | 10.72% |
| % background sequences | 4.65% |

**Extended Data Figure 3.** Inducible BAF complex binding over FOS binding sites. For all boxplots: line = median, box = IQR (Q1-Q3), whiskers = 1.5 x IQR, dots = outliers. (**a**) Boxplot of the $\log_2$ BAF complex subunit CUT&Tag signal normalized to paired IgG control +1 in H1- and HUES64-derived hpiNs over H9-derived hpiN FOS binding sites, before (blue) or 2 hr after (red) depolarization (HUES64 n = 3 biological replicates, H1 ARID1A = 2 biological replicates, H1 ARID1B = 4 biological replicates, **p = 1.35e-13, *** p<2.2e-16 Wilcoxon signed rank test). (**b**) Scatter plot of sequencing reads (unstimulated vs 2 hr post-depolarization) for all NPAS4 binding sites. Sites with significantly different NPAS4 binding between timepoints are highlighted in red (n = 5 biological replicates, adjusted p<0.05). (**c**) Boxplot of the $\log_2$ ratio +1 of BAF complex subunit CUT&Tag normalized signal in stimulated versus unstimulated hpiNs over FOS (black) or NPAS4 (green) binding sites (n = 3, ***p<2.2e-16 Wilcoxon unpaired test). (**d**) Enrichment for the binding motif for Arnt, a known NPAS4 interactor, in NPAS4 binding sites. (**e**) Paired open chromatin regions (n = 264 pairs) categorized by high (black) or low (pink) stimulated/unstimulated ratios of FOS CUT&Tag normalized signal (left panel) but similar stimulated/unstimulated ratios of ATAC-seq signal (right panel) (***p<2.2e-16, Wilcoxon signed rank test). (**f**) Boxplot of the $\log_2$ stimulated/unstimulated ratio +1 of BAF complex subunit CUT&Tag normalized signal in regions with high versus low FOS ratios. (**g**) EBF motif enrichment in FOS/BAF binding sites.

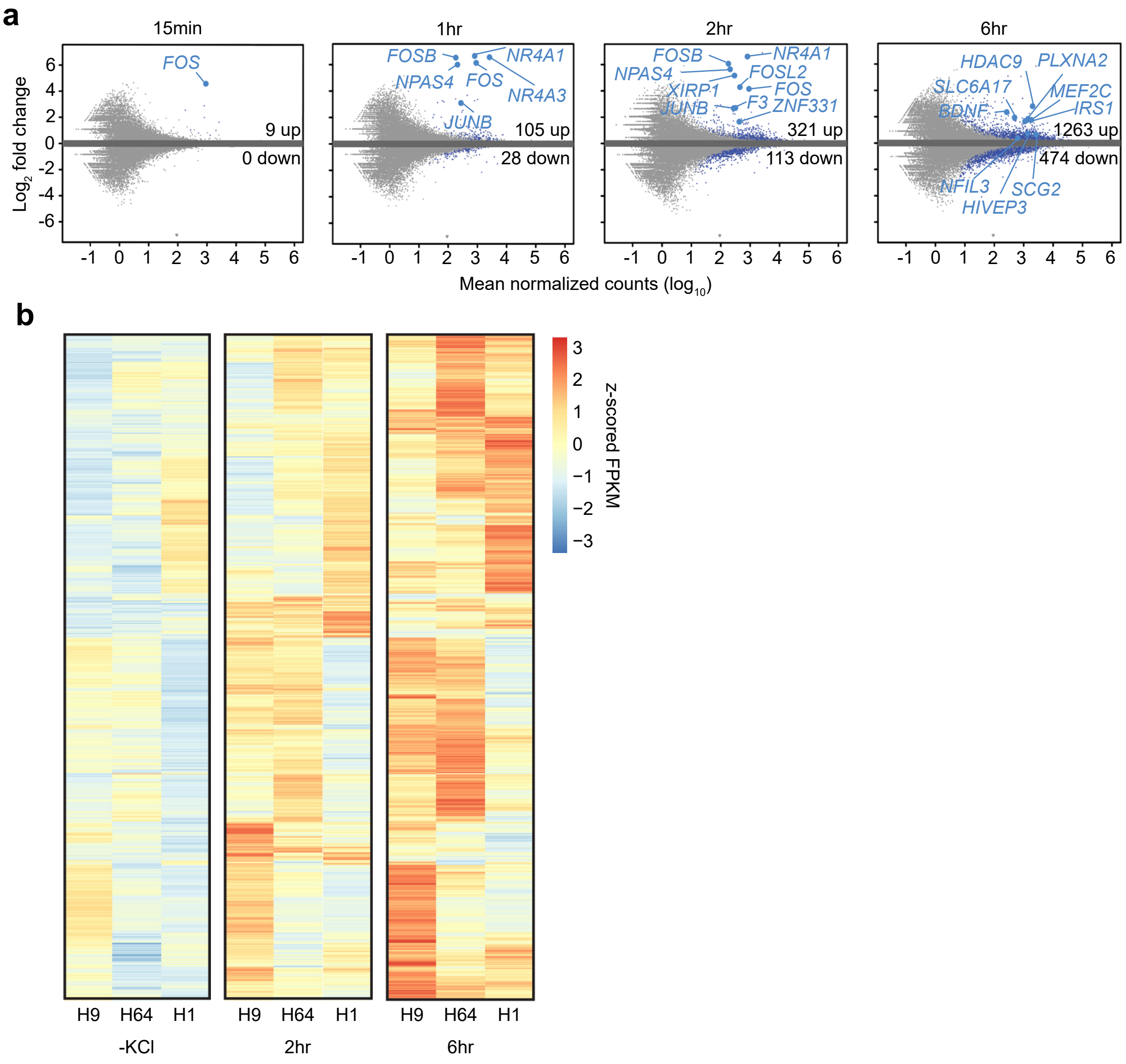


**Extended Data Figure 4.** Activity-dependent gene expression in hpiNs. (**a**) MA plots of RNAseq data comparing unstimulated hpiNs to stimulated hpiNs. Differentially expressed genes highlighted in blue (n = 3 biological replicates except for the 2 hr timepoint which had 2 biological replicates, adjusted p<0.05). (**b**) Heatmap representing RNA expression (z-scored FPKM) of H9-derived hpiN activity-dependent genes across hpiNs derived from H9, H1, or HUES64 hESCs (n = 3 biological replicates per line).

**a** **b**

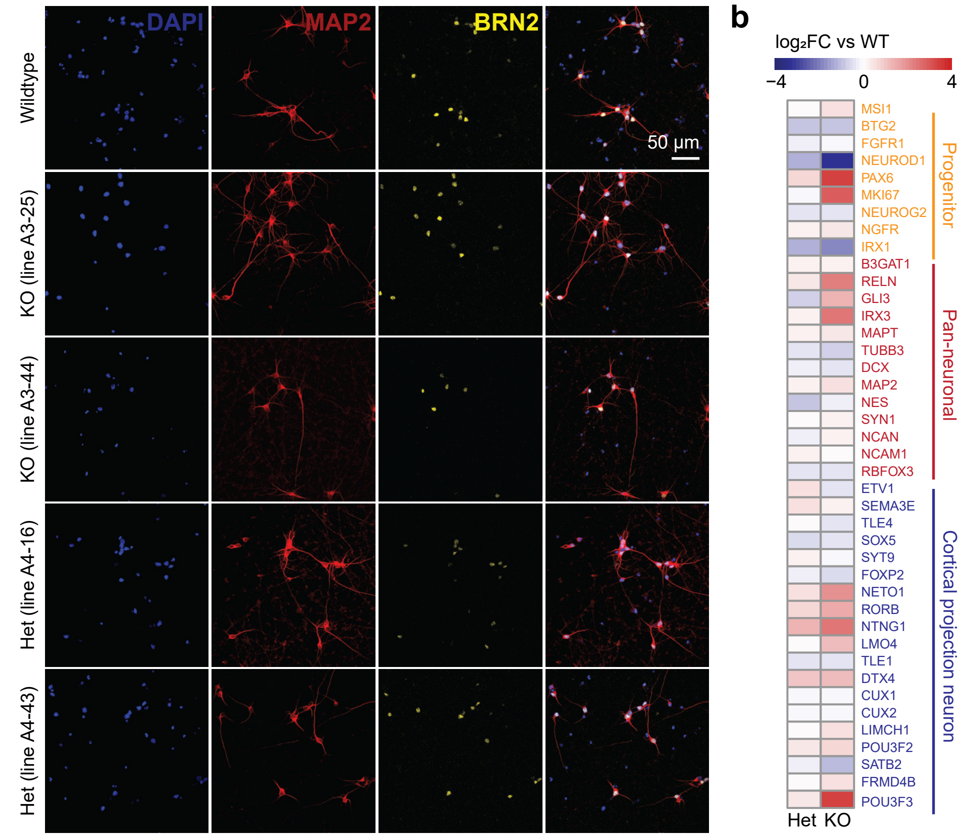


**c**

| | -KCl dec v WT | -KCl inc v WT | 2hr dec v WT | 2hr inc v WT | 6hr dec v WT | 6hr inc v WT |
|---|---|---|---|---|---|---|
| Het | 1322 | 388 | 604 | 435 | 336 | 459 |
| KO | 10494 | 1680 | 10627 | 2156 | 7477 | 1873 |

**d**

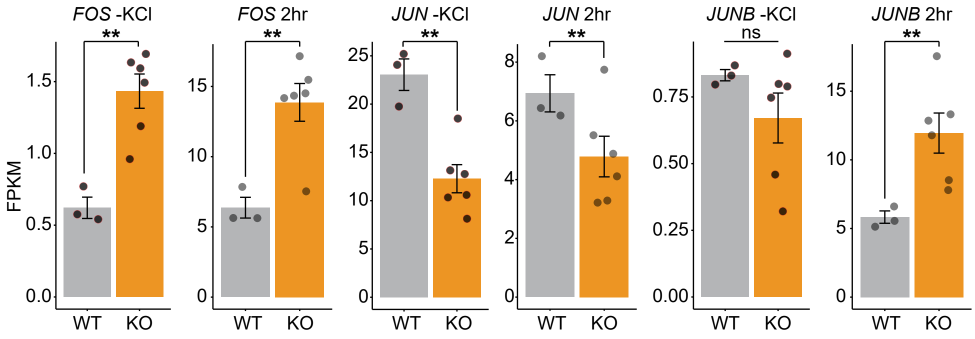


**Extended Data Figure 5.** Characterization of ARID1A het and KO hpiNs. (**a**) Immunohistochemistry for the neuronal markers MAP2 and BRN2 in wildtype, ARID1A KO, and ARID1A het hpiNs at Day 28. (**b**) Heatmap showing the $\log_2$ fold-change in marker gene RNA expression in ARID1A het or KO hpiNs compared to wildtype hpiNs at Day 28. (**c**) Table of regions exhibiting differential chromatin accessibility identified by ATAC-seq in ARID1A het and KO hpiNs compared to wildtype hpiN cultures (adjusted $p<0.05$, $|\log_2FC|>1$). Dec = decreased in mutant line compared to wildtype; inc = increased in mutant line compared to wildtype. (**d**) Bar graphs showing *FOS*, *JUN,* or *JUNB* RNA expression (FPKM) before (-KCl, blue) or 2 hr after (red) depolarization in WT and ARID1A KO (KO) hpiNs (n = 3 biological replicates, mean ± s.em., dots = individual data points).

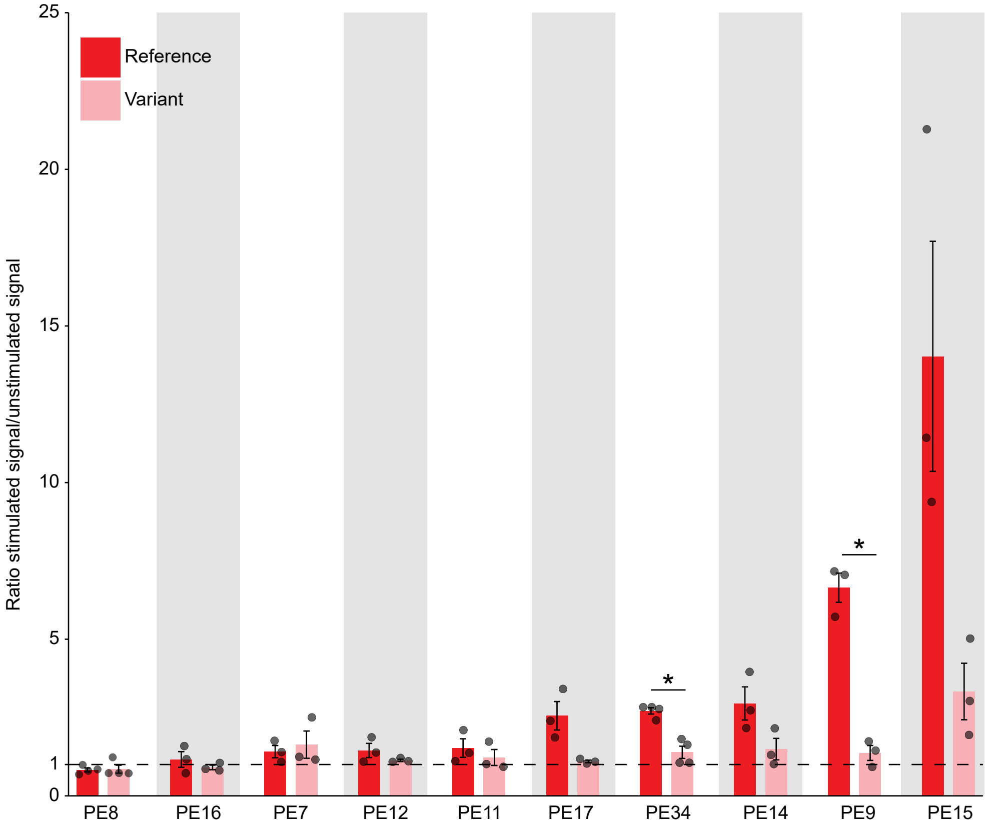


**Extended Data Figure 6.** Impact of SNVs in the AP-1 motif on activity-dependent enhancer function. Bar graph showing the ratio of depolarized versus unstimulated reporter assay signal driven by 500 bp enhancer regions with the reference sequence (dark red) compared to the same 500 bp regions with a single SNV introduced in the AP-1 motif (light red) for all tested PEs (mean ± s.e.m., dots = mean of 2 technical replicates for each biological replicate, n = 4 biological replicates for PE8 and PE34, n = 3 biological replicates for all other PEs, *adjusted $p<0.05$). Cultured mouse E16.5 cortical neurons were transfected on Day 3-4, silenced on Day 5, and depolarized with addition of KCl on Day 6.
